# The Walking Brain: Mapping feature-specific cortical responses to naturally unfolding sensory experience in the real world

**DOI:** 10.64898/2026.09.29.754498

**Authors:** Miray Altınkaynak, Laura B. Carlton, Ege Topkoç, David A. Boas, Meryem A. Yücel

## Abstract

Understanding how the brain processes the everyday world is fundamental, yet laboratory paradigms exclude self-generated movement, unscripted events, and a continuously evolving multisensory environment. Wearable neuroimaging enables real-world measurement, but linking neural activity to uncontrolled sensory information remains challenging. Here, we extend a feature-based framework from naturalistic movie viewing to everyday behavior using wearable whole-head high-density fNIRS and participant-specific audiovisual recordings during movies and outdoor walking with no explicit task or scripted audiovisual sequence. Luminance-and sound-related cortical maps showed significant spatial correspondence, particularly in visual and auditory regions, whereas face-related correspondence was weaker. Luminance-and face-related maps were reproducible within individuals across repeated walks. Additional exploratory analysis demonstrated, to our knowledge, the first out-of-sample encoding of continuous environmental features during unconstrained real-world fNIRS, predicting cortical activity during held-out walking periods. Together, these findings establish a feature-based framework for linking cortical activity to naturally unfolding sensory experience in everyday behavior.

## Introduction

Naturalistic neuroscience has transformed the study of human brain function by replacing highly simplified stimuli with movies, narratives and other continuous experiences that more closely resemble everyday life^1–8^. These paradigms have revealed robust and reproducible patterns of activity across distributed cortical systems^6,9–14^ and have enabled continuous properties of visual and auditory experience to be related directly to ongoing brain activity, yielding feature-specific patterns of cortical organization^15,16^. Extending this feature-based approach to real-world behavior, however, presents a fundamental challenge: outside the laboratory, the sensory stream is no longer predefined by the experimenter but unfolds continuously as individuals move through and interact with their environment. Whether feature-specific cortical organization can be recovered under these conditions, and how the resulting maps relate to those obtained in the laboratory, remains unknown. Addressing these questions is important for developing principled approaches to interpreting brain activity during naturally unfolding behavior.

Even highly naturalistic laboratory paradigms retain substantial experimental structure. Participants are typically stationary and visual attention is strongly influenced by cinematography, narrative structure or task instructions^9,17^. In everyday experience, by contrast, visual sampling is self-directed and sensory input changes continuously with movement^3–5^. This creates both a scientific and an analytical challenge. Conventional neuroimaging experiments begin with a known stimulus or task structure and ask how the brain responds to it. In unstructured real-world behavior, the relevant experimental variables must instead be recovered from the participant’s unfolding experience. Recording and quantifying that experience therefore offers a possible route toward studying brain function without imposing an explicit cognitive task or scripted sequence of sensory events.

Feature-based encoding provides a natural framework for bridging these two settings because it establishes quantitative relationships between continuously varying properties of experience and ongoing brain activity. Pioneering fMRI work demonstrated that models trained on natural movies could predict cortical responses to novel movie stimuli and support reconstruction of viewed visual experience^12^. Related encoding studies subsequently revealed distributed representations of semantic and other complex properties of natural stimuli^18–21^. This framework has recently begun to extend to high-density optical neuroimaging. High-density diffuse optical tomography (HD-DOT) studies demonstrated feature-specific cortical mapping during naturalistic movie viewing^22^, subsequent work showed that model-based representations could identify novel movie clips outside the training set^23^, and more recent high-dimensional semantic encoding models have predicted HD-DOT responses during movie viewing and supported identification of viewed clips^24^. Together, these studies show that distributed information contained in naturalistic stimuli can be recovered using optical measurements. Critically, however, the sensory stream in these studies remained experimenter-defined. Whether feature-based approaches can recover interpretable cortical organization when sensory information is encountered naturally, rather than experimentally prescribed, remains unknown.

Addressing this question requires distributed cortical measurements that can accompany natural behavior. Mobile EEG and wearable fNIRS have increasingly enabled brain imaging beyond the laboratory^3,4,25^, including studies of navigation, walking, memory and cognitive workload^26–32^. Yet most real-world neuroimaging studies retain substantial experimental structure, for example by requiring predefined cognitive tasks, manipulating specific events or targeting restricted cortical regions^28–35^. Thus, an important gap remains between taking a conventional experiment into the real world and measuring cortical organization during behavior in which the moment-to-moment sensory sequence itself unfolds naturally. Wearable whole-head high-density fNIRS offers an opportunity to narrow this gap by providing distributed cortical coverage during movement while simultaneously recording the participant-specific environment from which continuous sensory variables can be reconstructed.

Here, we combined wearable whole-head high-density fNIRS, mobile eye tracking and participant-specific scene-camera recordings to extend feature-based cortical mapping to natural outdoor behavior. Participants followed a predefined outdoor route but performed no explicit cognitive task and experienced no experimentally scripted audiovisual sequence. Instead, their visual and auditory input varied naturally as they moved through the environment, and the resulting sensory stream was reconstructed from their recordings. We used movie viewing as a controlled naturalistic benchmark and extracted analogous continuous audiovisual features from the movie and from each participant’s outdoor experience. This allowed us to ask whether feature-specific cortical maps recovered during outdoor walking corresponded spatially to those measured during movie viewing, and whether real-world maps were reproducible across repeated walks despite changes in the events encountered. Together, these analyses provide, to our knowledge, the first demonstration that feature-specific cortical organization can be recovered during natural real-world behavior, while also establishing a common framework for quantifying its spatial correspondence with cortical organization measured under controlled naturalistic conditions. More broadly, this framework provides a route from experiments in which the investigator defines the sensory events to neuroimaging in which those events are reconstructed from naturally unfolding experience. Finally, as an exploratory extension, we asked whether a broader multivariate representation of the outdoor sensory environment contained sufficient information to predict cortical hemodynamics during held-out periods of walking. To our knowledge, this is the first application of continuous environmental encoding to predict held-out fNIRS cortical dynamics during natural outdoor behavior.

## Results

### Feature-specific cortical response maps

We first characterized cortical responses to analogous sensory features encountered during controlled movie viewing and natural outdoor behavior. We recorded whole-head high-density fNIRS during naturalistic movie viewing in the laboratory (n = 18) and during two walking runs along a predefined outdoor route (n = 20), with participant-specific scene-camera recordings capturing the naturally unfolding audiovisual environment during outdoor walking (Fig.1).

**Fig. 1.**
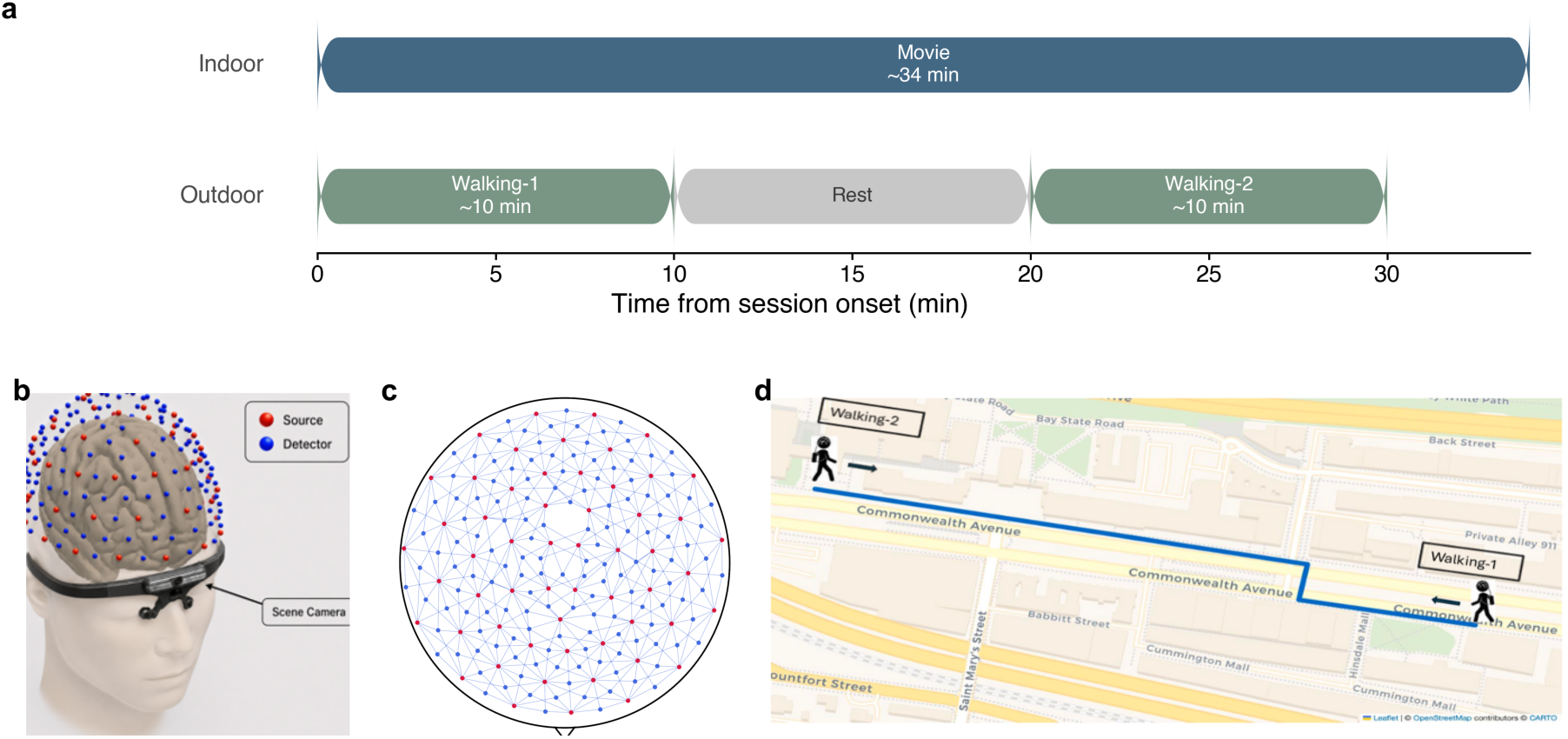
Experimental paradigm and setup. **a,** Timing of the two recording sessions. In the indoor session participants viewed a naturalistic movie (∼34 min). In the outdoor session, participants completed two walking runs along the same predefined route (∼10 min each), separated by a 10-min sitting recording used only in the prediction analysis. b, Three-dimensional layout of the whole-head HD-fNIRS system comprising 56 sources and 144 detectors (567 measurement channels), shown together with the scene camera of the head-mounted eye-tracking system. **c,** Two-dimensional probe layout; each blue line represents one measurement channel. First-nearest-neighbor source–detector separations are 19 mm and second-nearest-neighbor separations are 33 mm. **d,** Outdoor walking route. Participants walked at their own pace; the two runs followed the same route in opposite directions.

Continuous audiovisual features were extracted from the movie and outdoor recordings, convolved with a canonical hemodynamic response function (HRF), and incorporated into condition-specific design matrices (Fig. 2). fNIRS data from both environments were processed using the same analysis pipeline, comprising channel-space preprocessing, cortical image reconstruction, parcellation, and parcel-wise statistical modelling (Methods and Fig. 8).

**Fig. 2.**
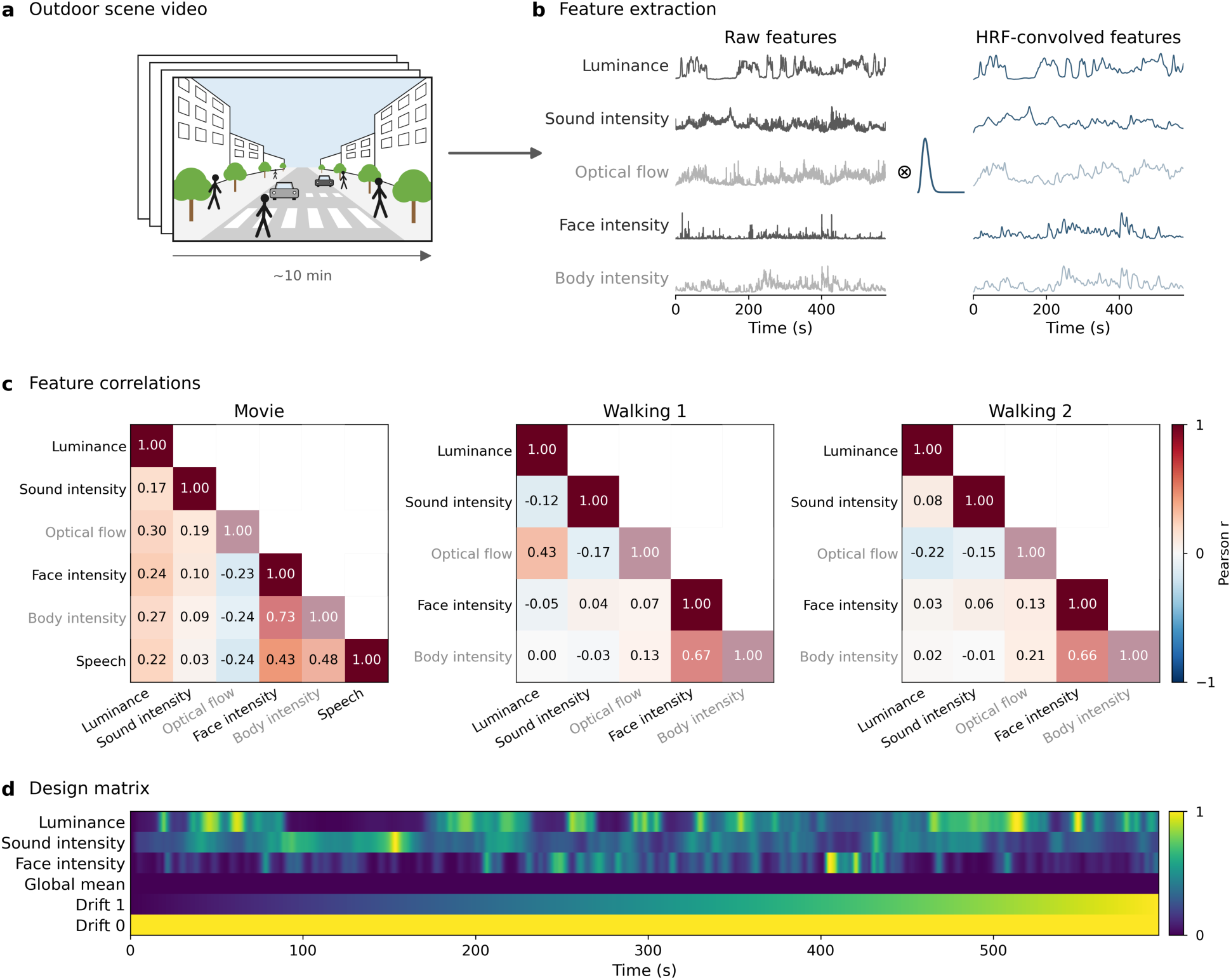
Feature extraction and design matrix construction. **a,** Schematic of the egocentric video recorded by the head-mounted scene camera during an outdoor walking run (∼10 min). **b,** Five features: luminance, sound intensity, optical flow, face intensity, and body intensity were extracted frame-by-frame from the recordings and convolved with a single-gamma HRF. Traces are scaled independently for display. Features excluded from the final model are shown in grey. **c,** Pearson correlations among HRF-convolved features for movie viewing and the two outdoor walking runs. Body intensity and optical flow were excluded from the final model because of their high correlations with face intensity and luminance, respectively (see Methods). **d,** Final design matrix, comprising the three retained features min–max normalized to the unit interval together with a global mean regressor and first-order drift terms; nuisance regressors are rescaled here for display only. The movie model additionally included speech presence.

Feature-specific group-level *t* maps were estimated independently for Movie, Walking 1, and Walking 2 using the same GLM framework and are displayed on a common color scale within each feature (Fig. 3).

**Fig. 3.**
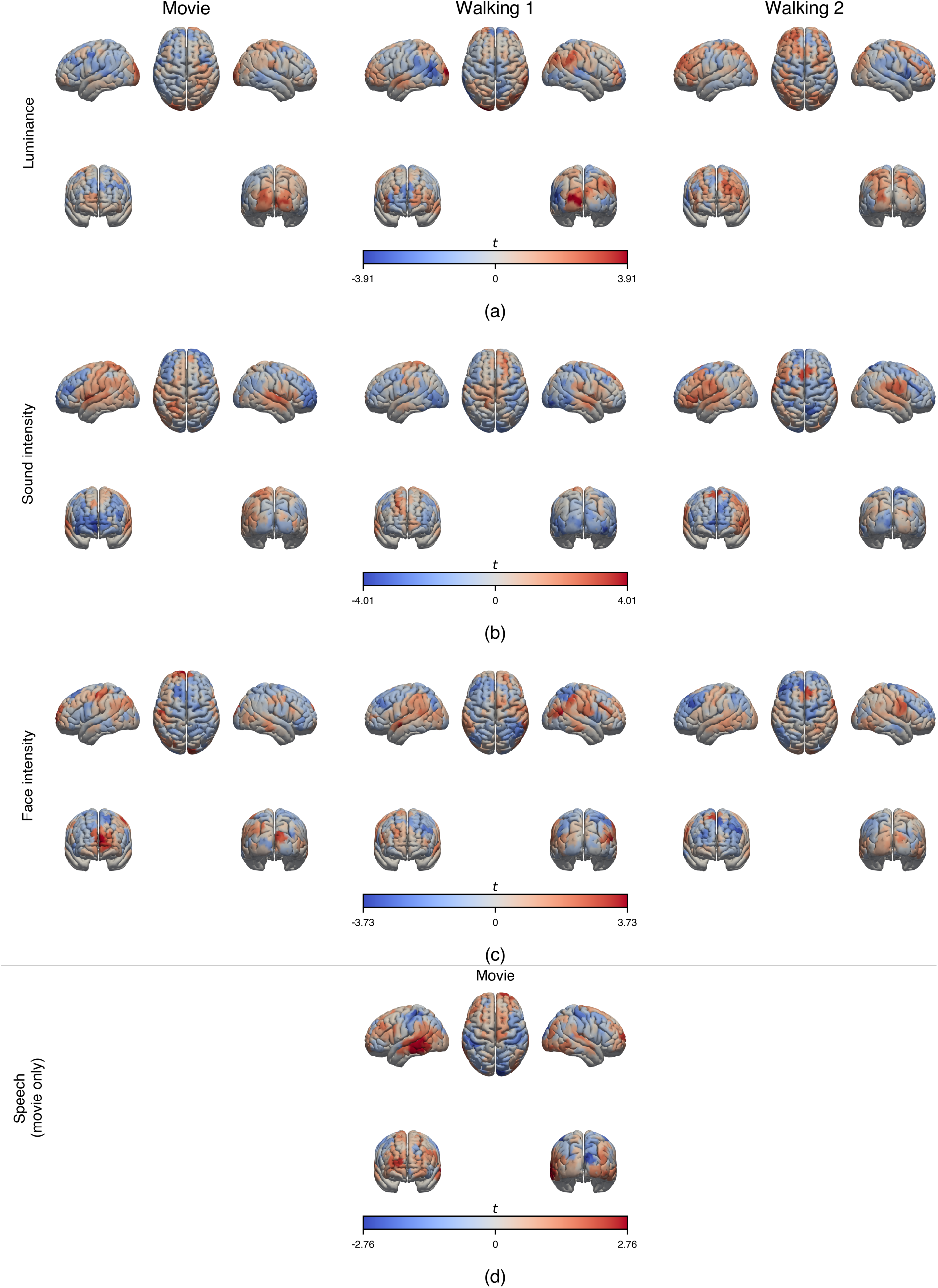
Feature-specific cortical response maps. Group-level *t* maps for HbO on the ICBM152 cortical surface, shown for **(a)** luminance, **(b)** sound intensity, **(c)** face intensity and **(d)** speech. Each block gives left lateral, superior and right lateral views (upper row) and anterior and posterior views (lower row). Color scales are set separately for each feature at 90% of its largest |*t*| (a: ±3.91; b: ±4.01; c: ±3.73; d: ±2.76) and are shared across the conditions within a row, so maps can be compared along a row. Maps are unthresholded. Speech was modelled for the movie only.

Luminance responses were most prominent over occipital cortex and appeared more left-lateralized during walking than during movie viewing. Sound intensity responses were concentrated over temporal regions, most clearly during movie viewing and Walking 2. Face-related responses appeared more spatially dispersed across conditions. Extrastriate visual cortex was the only region showing consistent face-related activation across all three sessions, indoor and outdoor alike. Speech responses, estimated for the movie only, elicited bilateral temporal cortical responses with a clear left-hemisphere predominance, extending beyond primary auditory cortex into superior and middle temporal association regions.

### Spatial correspondence of feature-specific maps across environments

We next assessed the spatial correspondence of feature-specific cortical response patterns across controlled movie viewing and natural outdoor behavior. To quantify spatial correspondence, we correlated the group-level t maps of each condition pair parcel by parcel, and assessed significance against a spin permutation null that preserves spatial autocorrelation (5,000 rotations; BH-FDR corrected; all values are reported in Supplementary Table 1).

In the whole-brain analysis (Fig. 4a), luminance maps showed significant spatial correspondence between the Movie and Walking 1 (r = 0.28, *P* = 0.001) and between the two walking runs (r = 0.29, *P* < 0.001), but not between the Movie and Walking 2 (r = 0.03, *P* = 0.74). For sound intensity, significant correspondence was observed between Movie and Walking 1 (r = 0.24, *P* = 0.046) and between Movie and Walking 2 (r = 0.32, *P* = 0.003), whereas the correspondence between the two walking runs was not significant (r = 0.06, *P* = 0.58). No comparison reached significance for face intensity (all *P* ≥ 0.25).

**Fig. 4.**
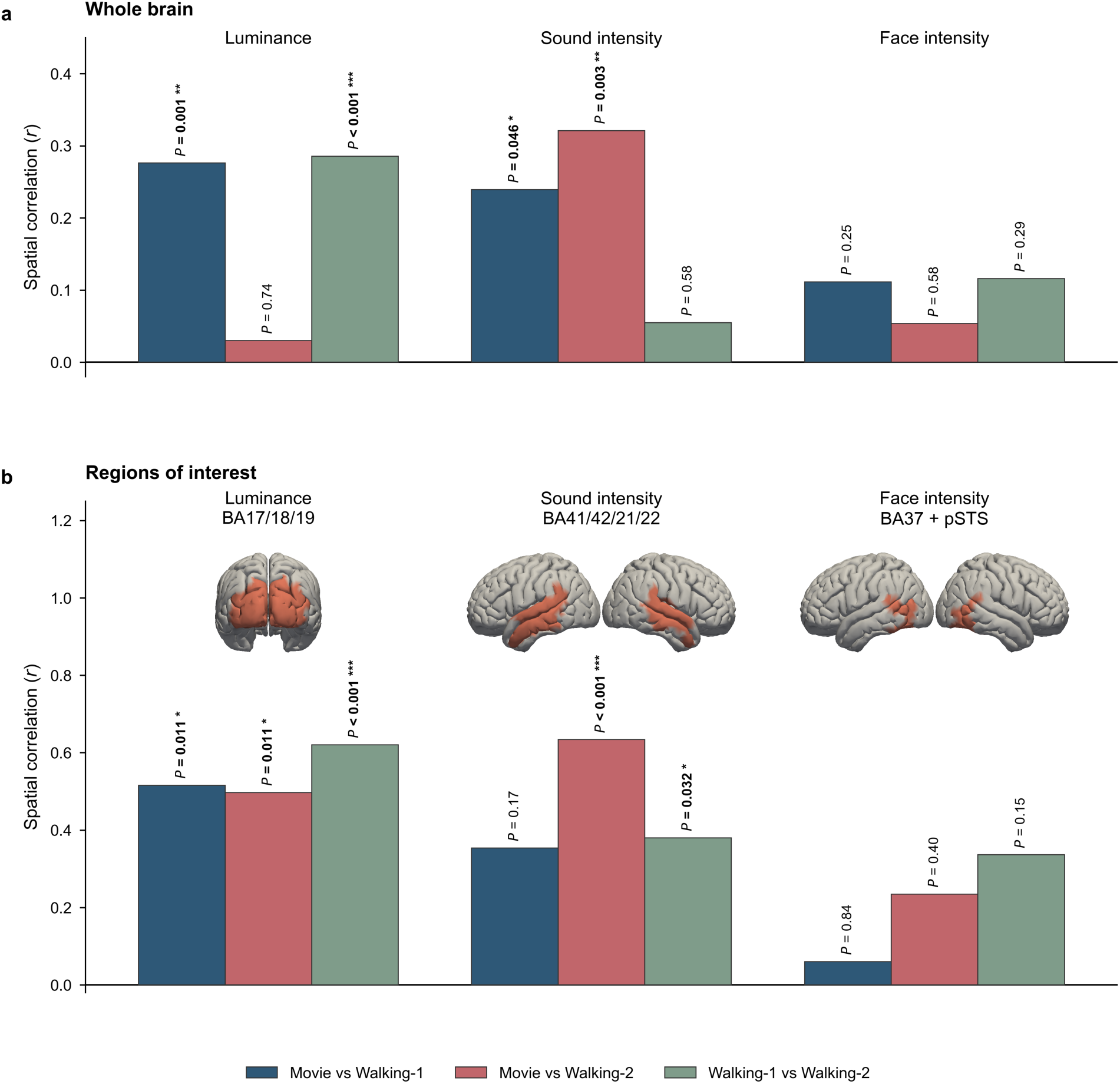
Spatial correspondence of feature-specific cortical response maps across conditions. Parcel-wise correlations between group-level HbO *t* maps for each pair of conditions, computed **a,** across the whole brain and **b,** within a priori regions of interest (ROIs; insets). Significance was assessed against spatial null distributions generated using spin permutations (5,000 rotations), which preserve the spatial autocorrelation structure of cortical maps. *P* values were BH-FDR corrected separately for the six movie–walking and three walking–walking comparisons within each analysis. Adjusted *P* values are shown above each bar; \**P* < 0.05, \*\**P* < 0.01, \*\*\**P* < 0.001.

In the ROI analysis (Fig. 4b), luminance showed significant spatial correspondence for all three condition pairs (r = 0.50–0.62, all *P* ≤ 0.011). Sound intensity was significant within the auditory ROI for Movie–Walking 2 (r = 0.63, *P* < 0.001) and between walking runs (r = 0.38, *P* = 0.032) but not for Movie–Walking 1 despite a moderate correlation (r = 0.35, *P* = 0.17). Face intensity maps did not show significant correspondence in either the whole-brain or ROI analyses, although the highest correlation was observed between Walking 1 and Walking 2 (r = 0.34, *P* = 0.15).

### Reproducibility of feature-specific cortical maps

Within-subject reproducibility was assessed at the whole-brain level by correlating parcel-wise *t* maps estimated from the first and second halves of the movie and, separately, from the two walking runs. Fisher z-transformed correlations were tested against zero across participants (sign-flip permutation, 5,000 permutations; BH-FDR across features within each dataset). For the movie (Fig. 5a), no feature showed significant within-subject reproducibility across the two halves (all FDR-adjusted *P* = 0.46; mean r between −0.05 and 0.09). For walking (Fig. 5b), luminance (mean r = 0.16, *P* = 0.011) and face intensity (mean r = 0.10, *P* = 0.007) showed significant within-subject reproducibility across runs, while sound intensity showed a similar but non-significant effect (mean r = 0.08, *P* = 0.071). At the individual level, positive Walking 1–Walking 2 correlations were observed in 15/20 participants for luminance, 14/20 for sound intensity, and 16/20 for face intensity (Fig. 5c).

**Fig. 5.**
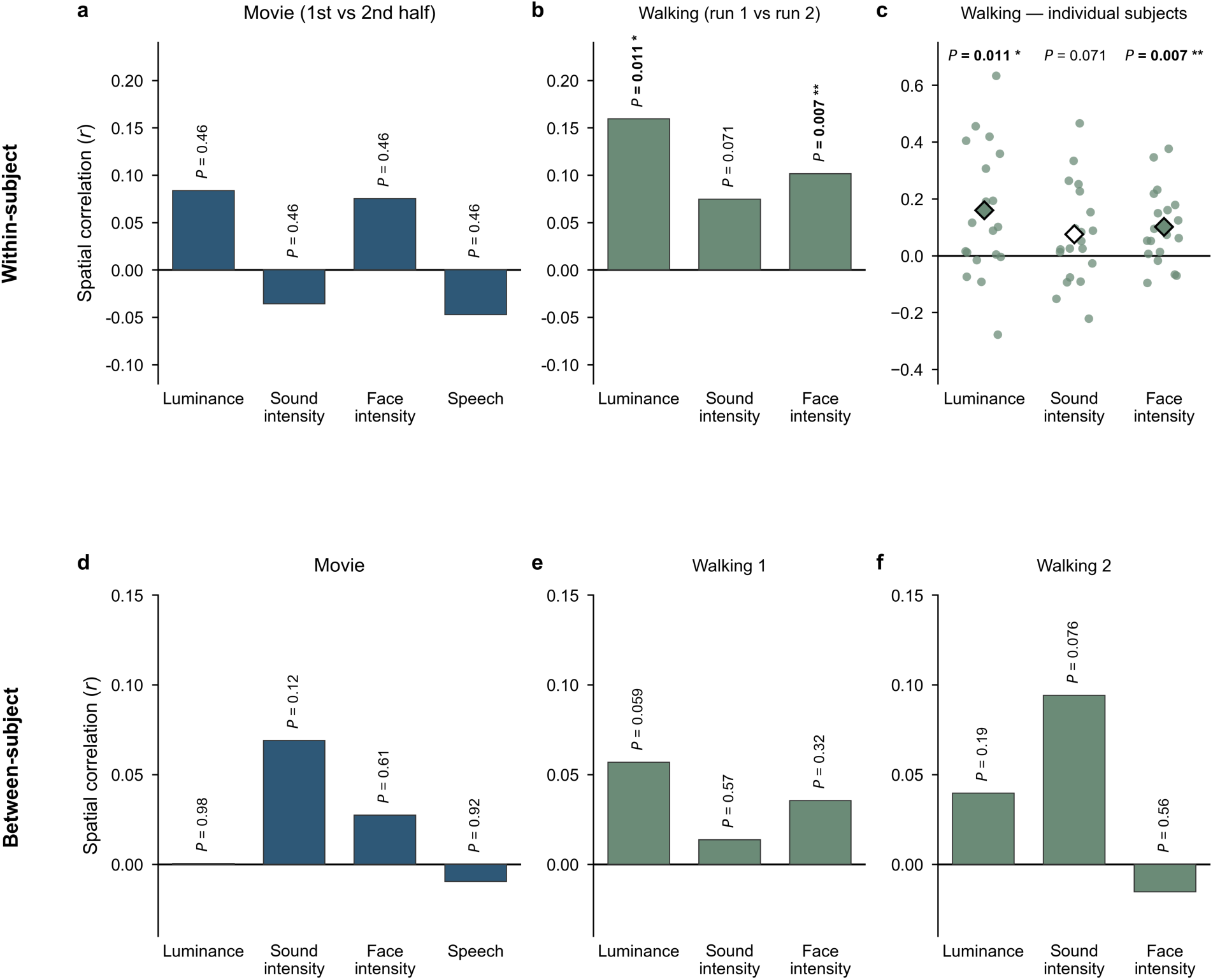
Reproducibility of feature-specific cortical response maps. **a–c**, Within-subject reproducibility. **a**, Split-half spatial correlations between feature-specific HbO *t* maps estimated from the first and second halves of the movie for each participant. **b**, Spatial correlations between feature-specific HbO *t* maps estimated from the two outdoor walking runs for each participant. **c**, Individual Walking 1–Walking 2 spatial correlations; each point represents one participant, and diamonds indicate the group mean. **d–f**, Between-subject reproducibility, quantified by spatially correlating each participant’s feature-specific *t* map with the corresponding group map estimated from the remaining participants (leave-one-out), for Movie (**d**), Walking 1 (**e**) and Walking 2 (**f**). Correlation coefficients were Fisher *z*-transformed and tested against zero across participants using sign-flip permutation tests (5,000 permutations), with BH-FDR correction across features within each analysis. FDR-adjusted *P* values are shown above each comparison; \**P* < 0.05, \*\**P* < 0.01, \*\*\**P* < 0.001. Speech was modelled for the movie only.

Between-subject reproducibility was assessed at the whole-brain level using a leave-one-out (LOO) approach, in which each participant’s feature-specific *t* map was spatially correlated with the corresponding group map estimated from the remaining participants. No feature showed significant between-subject reproducibility in any condition (Fig. 5d–f; all *P* ≥ 0.059) although mean correlations were positive in nearly all feature–condition combinations, with the largest mean correlations for luminance during Walking 1 and sound intensity during Walking 2 (Fig. 5e,f).

### Environmental features predicted unseen cortical dynamics during outdoor walking

We next asked whether environmental features could predict ongoing cortical hemodynamics during real-world behavior. Subject-specific ridge-encoding models used residual optical flow, environmental sound intensity, luminance, body intensity and face intensity, represented at delays of 1–12 s. Out-of-sample performance was evaluated using a nested blocked design (Fig. 6a). In each of four outer folds, one nonoverlapping half of a walking run was reserved for testing, whereas the sitting run and the remaining walking data formed the development set. Ridge regularization was selected by fivefold blocked cross-validation within the development data before the final model was refitted and evaluated once on the held-out walking segment.

**Fig.6.**
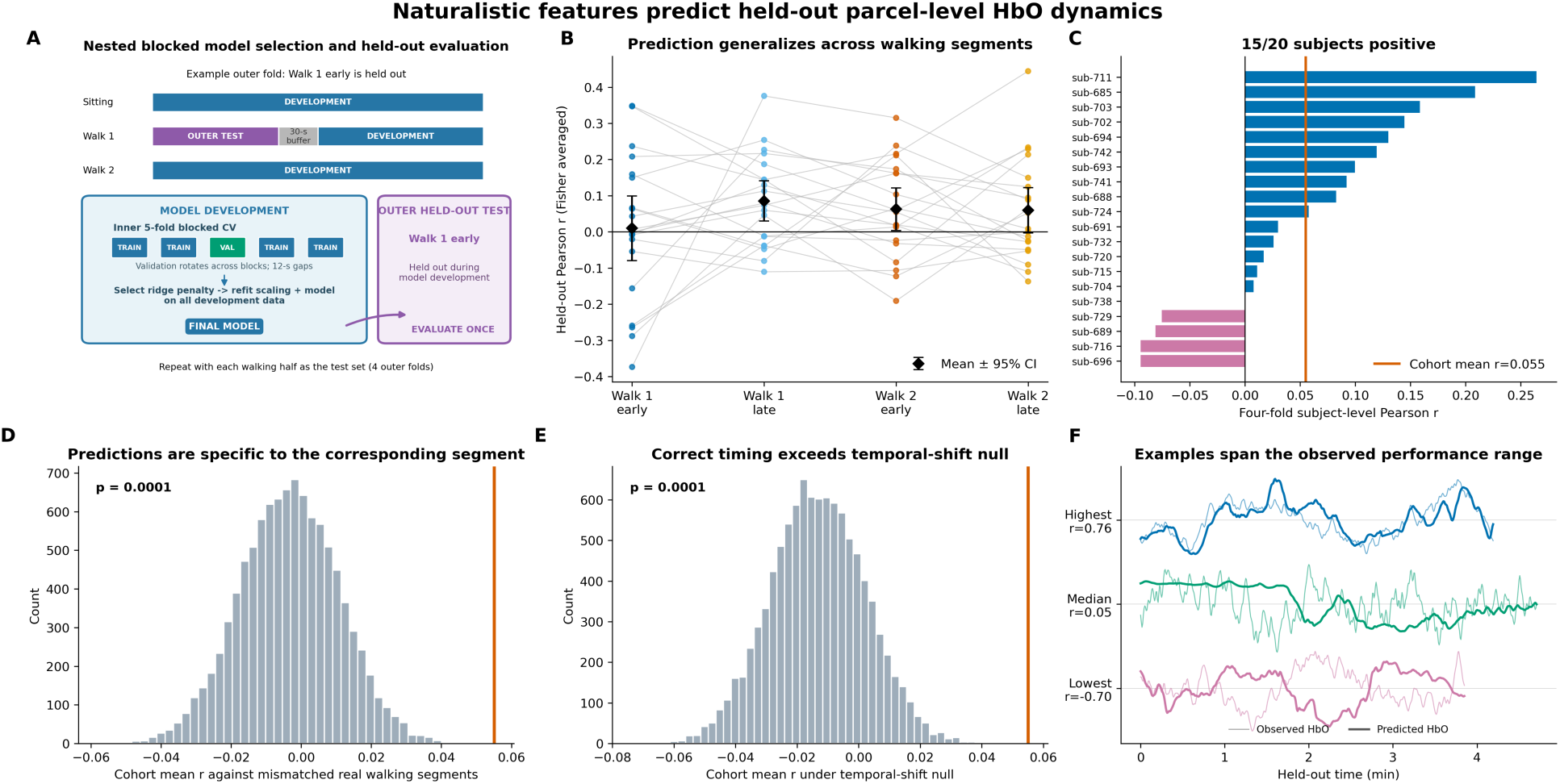
Naturalistic features predict held-out parcel-level HbO dynamics. **a,** Nested blocked evaluation design, illustrated for an outer fold in which the early half of walking run 1 was the untouched test set. The sitting run, walking run 2 and the nonheld-out portion of walking run 1 formed the development set; a 30-s buffer adjoining the test boundary was excluded. Fivefold contiguous blocked cross-validation within the development set, with 12-s gaps around validation blocks, selected the ridge penalty. Scaling and ridge coefficients were then refitted using all development data before predicting the untouched test segment once. The procedure was repeated with each early and late walking half serving as the test set, producing four outer folds. **b,** Generalization performance across the four nonoverlapping outer test folds. Each colored point is one participant’s held-out Pearson correlation, calculated parcel-wise and Fisher-*z* averaged within the indicated fold; gray lines connect folds from the same participant. Black diamonds and error bars show the across-participant mean and 95% confidence interval for each fold. **c,** Participant-level prediction accuracy after Fisher-*z* averaging parcel correlations across all four outer folds and transforming back to correlation units. Horizontal bars represent individual participants, ordered by performance; blue and magenta indicate positive and negative correlations. The orange line marks the cohort mean (r = 0.055); 15 of 20 participants had positive four-fold average correlations. The participant-bootstrap 95% confidence interval was 0.013–0.098 (100,000 resamples). **d,** Correct-segment specificity control. Each held-out prediction was paired with equal-length, noncorresponding windows drawn from the same participant’s other real walking data; candidate windows advanced in 60-s steps and sitting data were excluded. The histogram shows 10,000 cohort statistics, each sampling one mismatched window per participant and fold and repeating the complete Fisher-*z* aggregation. The orange line denotes the correctly corresponding prediction-HbO statistic (one-sided Monte Carlo *P* = 1 × 10⁻⁴, plus-one correction). **e,** Autocorrelation-preserving temporal-shift control. Held-out predictions were circularly shifted relative to observed HbO, excluding shifts with circular distance below 60 s. The histogram shows 10,000 complete four-fold cohort aggregations under incorrect temporal alignment; the orange line shows the correctly aligned statistic (one-sided Monte Carlo *P* = 1 × 10⁻⁴, plus-one correction). **f,** Illustrative five-minute held-out predictions spanning the observed subject-parcel-fold performance range. Thin, lighter traces show observed standardized HbO and thicker traces show model predictions in the same color; vertical offsets are used only for display. Examples were selected algorithmically as the maximum, median-nearest and minimum parcel-wise held-out correlations across all eligible participant-parcel-fold combinations. All correlations were evaluated exclusively on outer held-out data. n = 20 participants.

Prediction accuracy varied across participants and held-out segments but was positive at the cohort level (Fig. 6b,c). After Fisher averaging parcel-wise correlations across the four outer folds within each participant, the mean held-out Pearson correlation was 0.055 (participant-bootstrap 95% CI, 0.013–0.098), and 15 of 20 participants had positive average correlations. Mean performance across participants was 0.010 for Walking 1 early, 0.085 for Walking 1 late, 0.063 for Walking 2 early, and 0.060 for Walking 2 late, indicating appreciable variation across participants and recording segments. Illustrative held-out time courses spanning the observed performance range demonstrate the correspondence associated with high, median, and negative parcel-level correlations (Fig. 6f).

Two empirical-null analyses tested whether this performance could be explained by generic slow fluctuations rather than correspondence between the environmental features and the corresponding HbO segment. First, held-out predictions were compared with noncorresponding windows from the same participant’s other walking data. Correct prediction-HbO pairings exceeded the resulting mismatched-segment null distribution (10,000 Monte Carlo iterations; one-sided *P* < 0.001; null mean *r* = -0.003; Fig. 6d). Second, predictions were circularly shifted relative to observed HbO while preserving their temporal structure and excluding shifts within 60 s of the correct alignment. Correct temporal alignment also exceeded this autocorrelation-preserving temporal-shift null distribution (10,000 iterations; one-sided *P* < 0.001; null mean *r* = -0.013; Fig. 6e). Together, these controls indicate that the observed prediction was specific to the corresponding walking segment and its temporal alignment, rather than arising solely from the smoothness of the signals.

### Prediction accuracy was spatially heterogeneous across cortical networks

We next characterized the spatial distribution of held-out prediction using the 247 Schaefer parcels that satisfied the sensitivity and quality-control criteria in all 20 participants (Fig. 7a). Parcel correlations were Fisher averaged across the four outer folds within participant and subsequently across participants. Prediction accuracy was spatially heterogeneous, with positive and near-zero values distributed across the measured cortex. Parcels not meeting the complete-participant criterion were excluded from the group map, ensuring that every displayed parcel had identical participant support.

**Fig.7.**
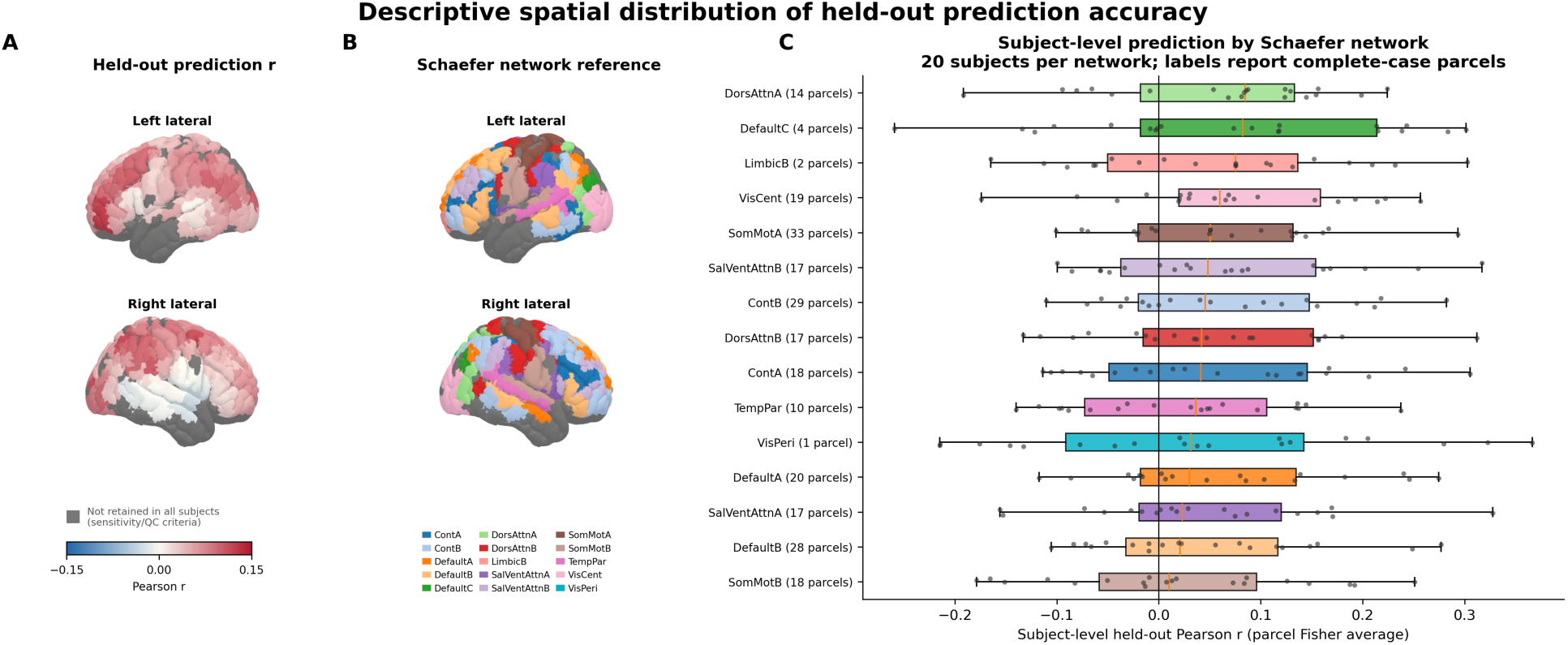
Descriptive spatial distribution of held-out prediction accuracy. **a,** Group descriptive map of parcel-wise held-out prediction accuracy. For each participant and Schaefer parcel, Pearson correlations between observed and predicted HbO were Fisher-z averaged across the four outer walking-half folds. Parcel values were subsequently Fisher-z averaged across all 20 participants and transformed back to correlation units. Only the 247 parcels satisfying sensitivity and quality-control criteria in every participant are colored. Parcels not retained in all participants are dark gray; off-white denotes an eligible parcel with correlation near zero. Blue and red indicate negative and positive correlations on a fixed scale of -0.15 to 0.15. **b,** Schaefer-network reference map for the same 247 complete-case parcels. Categorical colors identify the 15 Schaefer 17-network subdivisions represented by at least one universally retained parcel and match **c**; dark gray has the same exclusion meaning as in **a**. LimbicA and ContC are absent because neither contained a parcel retained in all 20 participants. **c,** Descriptive between-participant distributions of network-level held-out prediction. For each participant, parcel correlations within a network were Fisher-z averaged using the 247 complete-case parcels and transformed back to r, yielding one value per participant and network. Points show all 20 participants. Boxes show the median and interquartile range; whiskers extend to the most extreme observation within 1.5 times the interquartile range. Every network distribution contains 20 subject-level observations. Parenthetical counts in network labels report the number of complete-case parcels averaged within each participant, not participant count. Networks are ordered by median subject-level correlation and colors correspond to **b**. VisPeri contains one universally retained parcel but 20 subject-level correlations. Network summaries are descriptive; no network-specific hypothesis tests were performed. n = 20 participants.

For a network-level description, the 247 complete-case parcels represented 15 of the 17 Schaefer networks (Fig. 7b); LimbicA and ContC had no parcels retained in every participant. Within each participant, parcel correlations were Fisher averaged separately within each represented network, yielding 20 subject-level observations per network (Fig. 7c). The highest descriptive median correlations were observed in DorsAttnA (median *r* = 0.085; 14 parcels), DefaultC (median *r* = 0.083; 4 parcels), LimbicB (median *r* = 0.075; 2 parcels), and VisCent (median *r* = 0.060; 19 parcels). However, between-participant distributions overlapped substantially across networks and included negative values. These spatial results are therefore presented descriptively; no network-specific hypothesis tests or claims of significant differences between networks were made.

## Discussion

While naturalistic movie paradigms have provided a powerful framework for investigating feature-specific cortical representations under controlled laboratory conditions, whether feature-specific cortical organization can be recovered during natural real-world behavior remains largely unknown. Here, we extended this feature-based approach to natural outdoor behavior, estimating cortical maps from each participant’s naturally unfolding sensory experience, and compared them with maps obtained during laboratory movie viewing. Luminance-and sound-intensity-related cortical maps exhibited significant spatial correspondence across laboratory and outdoor environments, particularly within functionally relevant cortical regions, whereas face-related responses showed weaker correspondence. Furthermore, feature-specific maps obtained during outdoor walking showed significant reproducibility across two outdoor walking runs, indicating that individual maps contain a stable spatial component despite the challenges of unconstrained recording. Finally, an exploratory encoding analysis showed that continuously measured environmental features predicted cortical dynamics during held-out periods of outdoor walking, with prediction exceeding controls based on incorrect segment and temporal alignment. Together, these findings demonstrate that wearable whole-head HD-fNIRS enables the characterization of feature-specific cortical representations beyond controlled laboratory settings, providing a framework for investigating brain function during natural behavior.

Luminance and sound intensity are low-level sensory features that are reliably encoded during naturalistic audiovisual stimulation, driving robust and temporally consistent responses in early visual and auditory cortices^6,12^. Our findings extend these observations in a direction that could not previously be tested: the spatial organization of these low-level sensory representations shows significant correspondence across laboratory movie viewing and real-world behavior. This is particularly notable given the substantial differences between the two settings, including uncontrolled illumination and acoustic conditions, active locomotion, and self-directed visual sampling during real-world behavior.

In contrast, face-related cortical maps showed no significant correspondence across environments. One possible explanation is that participants likely attended to faces less frequently during outdoor walking than during movie viewing, even when faces were present in the scene. In addition, faces occurred in substantially different contexts across the two conditions: during movie viewing, they appeared within a structured narrative and cinematographically guided scenes, whereas during outdoor walking they belonged to real people encountered in a dynamic, self-directed environment.

### Luminance

Luminance is one of the fundamental low-level visual properties of natural scenes and has been widely used to characterize visual input. Previous studies have demonstrated that luminance strongly modulates responses in early visual cortex and that luminance information is represented throughout V1–V3, highlighting its fundamental role in cortical visual processing^36,37^. In agreement with these findings, luminance-related responses in our study were most prominent over occipital cortex. Interestingly, these responses were bilateral during movie viewing but showed a more left-lateralized occipital distribution during real-world walking.

Luminance maps showed significant correspondence across environments in most comparisons, both within the visual ROI and at the whole-brain level and were reproducible across repeated walking sessions. These results indicate that luminance-related cortical maps share a significant spatial component across laboratory and real-world conditions.

### Sound Intensity

Previous studies have shown that auditory responses scale with sound intensity or perceived loudness, highlighting acoustic energy as one of the primary drivers of early auditory cortical activity^38^. Naturalistic movie studies have extended these findings by demonstrating that continuous fluctuations in soundtrack intensity explain a substantial proportion of auditory cortical responses and that loudness is among the most robustly decodable auditory features^39^. Together, these findings establish sound intensity as a reliable feature for characterizing cortical auditory processing during naturalistic stimulation.

Consistent with this literature, sound intensity-related responses in our study were most prominent over bilateral temporal regions. At the whole-brain level, sound-intensity maps showed significant correspondence for both movie–walking comparisons. Within the auditory ROI, correspondence was significant between Movie and Walking 2 and between the two walking runs. The Movie– Walking 1 correlation within the auditory ROI was moderate (r = 0.35) but did not reach significance (P = 0.17). Together, these findings indicate that sound-intensity-related cortical maps share a significant spatial component across laboratory and real-world environments.

### Faces

Naturalistic movie paradigms have demonstrated that dynamic face perception recruits broader and more distributed cortical responses than static face viewing^40^. Here, we extended this comparison by contrasting face-related cortical maps obtained during movie viewing with those obtained during real-world outdoor walking. No significant cross-environment correspondence was observed for face-related maps, either within the face ROI or at the whole-brain level. During indoor viewing, face-related responses were predominantly localized to the medial prefrontal region, whereas outdoor walking was characterized by responses in sensorimotor and attention-related regions and this pattern was stable across both outdoor runs. Extrastriate visual cortex was the only region showing consistent face-related activation across all three sessions, indoor and outdoor alike.

The medial prefrontal response observed during movie viewing may reflect the embedding of faces within socially and narratively meaningful scenes rather than face processing per se. Consistent with this interpretation, naturalistic movie studies have implicated medial prefrontal cortex in processing social interactions^41^, character-related information^42^, and higher-order narrative context^43^. By contrast, faces encountered during outdoor walking occurred within an active behavioral context involving self-directed attention, locomotion, and the presence of real people in the surrounding environment, which may have contributed to the broader sensorimotor and attention-related pattern observed outdoors. Consistent with this possibility, a hyperscanning fNIRS study showed that live faces recruited stronger dorsal parietal responses and greater inter-brain coupling than the same faces viewed on a screen despite closely matched visual input^44^.

Together, these findings suggest that face-related cortical organization may be more sensitive to behavioral and social context than that associated with lower-level sensory features, potentially contributing to the absence of spatial correspondence across laboratory and real-world environments.

### Speech

Speech was modelled alongside sound intensity during movie viewing to distinguish speech-related responses from those associated with overall acoustic energy. The speech regressor elicited bilateral temporal responses with a clear left-hemisphere predominance, extending beyond primary auditory cortex into superior and middle temporal association regions, consistent with the distributed cortical organization of speech perception^45,46^.

Speech encountered during outdoor walking was sparse and incidental, arising primarily from passers-by rather than forming a sustained component of the auditory environment, and was therefore not modelled. Consequently, the spatial correspondence of speech-related cortical maps across laboratory and real-world environments could not be assessed in the present study. Future studies incorporating participant-specific speech models during real-world behavior will be important for determining how speech-related cortical organization compares across controlled and natural environments.

### Reproducibility of feature-specific cortical representations

Previous naturalistic movie studies have demonstrated reliable within-and between-subject reproducibility of brain responses using approaches such as voxel-wise inter-subject correlation (ISC)^6,22,47^, functional connectivity^48^, and connectivity-based functional topographies^49^, particularly within visual and auditory cortices. Importantly, this reproducibility has been shown to depend strongly on stimulus content, with meaningful and emotionally engaging movie content producing greater synchronization^50^. These studies, however, have primarily quantified reproducibility in the temporal domain, although spatial analogues of ISC have also been developed to assess the consistency of distributed response patterns across individuals^51,52^. Temporal measures of reproducibility, however, rely on repeated or shared stimulus sequences that provide a common temporal reference across recordings. Such temporal correspondence is inherently absent during unconstrained real-world behavior, where the sequence and timing of naturally encountered events differ across individuals and repeated walks. We therefore assessed reproducibility in the spatial domain, asking whether independently estimated cortical response maps for the same sensory feature retain a consistent spatial organization despite differences in the specific events encountered and their timing.

### Within-subject reproducibility

Previous studies have assessed within-subject reproducibility by having participants view the same movie repeatedly^22,53^. Because participants in the present study viewed the movie only once, a conventional test–retest comparison was not possible. Instead, we divided the 34-min movie into two 17-min segments and compared feature-specific cortical response maps derived from each segment within each participant. No significant split-half reproducibility was observed (Fig. 5a). Although our comparison was based on feature-specific cortical maps rather than the movie content itself, the two halves differed substantially in both feature distributions and narrative context. This content variability may have reduced the similarity of the resulting cortical maps (Supplementary Fig. 2a). Consistent with this interpretation, previous work has shown that both within-and between-subject reproducibility during movie viewing depends strongly on movie segment and emotional content^50^. Future studies using repeated presentations of the identical movie will be important for determining the test–retest reproducibility of feature-specific cortical representations.

Unlike the split-half movie comparison, the outdoor paradigm provided a closer approximation to a conventional test–retest design because participants completed two repeated walking sessions along the same route (Supplementary Fig. 2b). Luminance-and face-related cortical response maps showed significant within-subject reproducibility across the two walks, whereas sound intensity was positive but did not reach significance after FDR correction (Fig. 5b).

This finding is particularly noteworthy given the inherent variability of real-world environments: the specific events encountered differed between the two walks, yet the modelled features spanned comparable ranges. Feature-specific cortical response maps remained reproducible within individuals, demonstrating that meaningful cortical representations can be recovered even during unconstrained real-world behavior.

### Between-subject reproducibility

No significant between-subject reproducibility of feature-specific cortical response maps was observed during either movie viewing or outdoor walking. During movie viewing, this occurred despite all participants viewing the same naturalistic stimulus. To relate these findings to the conventional naturalistic movie literature, we additionally quantified parcel-wise temporal ISC in the same dataset (LOO; sign-flip permutation, 5,000 permutations; FDR-corrected). Temporal synchronization was significant across the sensitivity-masked cortex (r = 0.048, *P* < 0.001) and within both the visual (r = 0.060, *P* < 0.001) and auditory ROIs (r = 0.044, *P* = 0.001; Supplementary Fig. 3. and Supplementary Table 2), consistent with previous naturalistic studies^6,47,54^.

Unlike ISC, however, which quantifies the similarity of response time courses, our analysis compared the spatial organization of feature-specific response maps as described above. The presence of significant temporal synchronization in the same recordings therefore indicates that the absence of between-subject reproducibility of feature-specific maps does not reflect a lack of stimulus-locked neural responses. Between-subject comparison of feature-specific cortical maps is also limited by how precisely each individual map can be estimated. Detecting shared spatial organization across individuals will likely require longer recordings and more extensive sampling of each feature within participants, particularly given that maps estimated from single movie segments were not reproducible within participants (Fig. 5a).

During outdoor walking, additional sources of between-subject variability were present. Although all participants completed the same walking protocol, each individual experienced a unique sequence of visual and auditory events, sampled the environment differently through gaze, and likely allocated attention differently throughout the route. Attention may also at times have been directed toward internal processes not captured by the measured environmental features. Together, these sources of variability may limit the similarity of subject-specific feature maps across individuals.

Overall, although group-level maps showed significant spatial correspondence across conditions, between-subject reproducibility remained limited. Together with the significant within-subject reproducibility observed during repeated outdoor walking, these findings indicate that feature-specific cortical maps can contain a reproducible within-individual spatial component despite substantial variability across individuals. Future studies combining longer recordings with richer semantic feature spaces and participant-specific measures of visual sampling and attention may help determine whether more comprehensive characterization of each individual’s experience improves the detection of shared feature-related cortical organization across participants.

### Environmental features predict cortical dynamics during real-world walking

The encoding analysis provides initial evidence that a small set of continuously measured environmental features contains reproducible information about ongoing cortical hemodynamics during unconstrained outdoor walking. Although the mean held-out correlation was modest, prediction generalized to walking periods that were excluded from model selection and fitting, and most participants showed positive average performance. Modest effect sizes are expected in this setting: the measured features captured only a small part of the participant’s sensory environment, whereas cortical activity also reflects unmeasured scene content, behavior, internal thought, emotion, physiological state and measurement noise. In this context, the principal result is not that the model explains a large proportion of brain activity, but that even five relatively broad features predict a detectable component of previously unseen HbO dynamics outside the laboratory. The magnitude of the observed prediction can be considered in the context of previous high-density optical encoding studies. Fehner et al.^24^, for example, reported a mean held-out correlation of *r* = 0.032 during controlled naturalistic movie viewing using approximately 3.5 h of data per participant and a high-dimensional semantic feature space comprising 1,708 categories. Here, a mean held-out correlation of *r* = 0.055 was obtained from approximately 30 min of recorded data per participant during the outdoor session, using a small set of relatively basic sensory and environmental features. Although prediction performance cannot be directly compared across studies, this provides useful context for the magnitude observed here given the limited recording duration and unstructured nature of the recordings.

The two empirical-null controls support the specificity of the observed prediction. Correctly aligned predictions outperformed both circularly shifted versions of the same smooth time courses and pairings with noncorresponding real walking segments from the same participants. The temporal-shift analysis preserved the autocorrelation and amplitude distribution of the signals while disrupting their timing, whereas the mismatched-segment analysis retained genuine participant-specific outdoor HbO fluctuations while disrupting segment identity. Together, these controls make it less likely that the result reflects only slow hemodynamic structure, generic characteristics of walking or chance correspondence between smooth signals. They do not, however, establish a causal effect of any individual environmental feature, because the predictors covary naturally and the ridge model distributes information among correlated features and delays.

Prediction accuracy was spatially heterogeneous. Relatively higher descriptive subject-level medians were observed in dorsal-attention, central-visual and selected default-network subdivisions. Stronger prediction in dorsal-attention and visual cortex is broadly compatible with the continuous visuospatial orienting, optic flow and monitoring of people required during outdoor walking. The DefaultC pattern may reflect broader contextual changes that covary with the low-level, auditory and social features, but this interpretation is necessarily more tentative. Network distributions overlapped substantially, and some networks were represented by relatively few complete-case parcels. Accordingly, these maps should be viewed as anatomical summaries rather than evidence for significant differences among networks. The complete-case restriction ensured identical participant support for every displayed parcel, but it also reduced cortical coverage and preferentially excluded parcels that were small or inconsistently measurable after channel-level quality control.

The ridge-penalty sensitivity analysis indicated that the prediction result was stable over a wide range of strong regularization. This stability reduces concern that the reported performance depended on a particular upper grid boundary, but the absolute penalty magnitude and raw ridge coefficients should not be interpreted biologically. Ridge weights depend on scaling, sample count and correlations among delayed predictors, and Pearson correlation evaluates temporal correspondence rather than amplitude calibration. For this reason, the present study emphasizes held-out prediction rather than coefficient maps or feature-specific localization. An important next step will be to evaluate richer, prespecified scene representations in new data and determine whether they improve prediction while preserving strict separation between model development and final testing.

Overall, the encoding analysis provides a complementary test of whether environmental information measured during unstructured behavior contains temporally specific information about ongoing cortical activity. Beyond the spatial organization captured by the feature-mapping analyses, continuously measured environmental features predicted HbO dynamics during held-out periods of outdoor walking. Together, the mapping, reproducibility and prediction results demonstrate that feature-based approaches can extract interpretable, reproducible and predictive information from brain activity during naturally unfolding experience.

### Toward whole-head functional mapping in real-world environments

A key methodological contribution of the present study is the use of wearable whole-head high-density fNIRS combined with image reconstruction and parcel-based cortical analysis to characterize distributed cortical activity during unconstrained real-world behavior. This framework enabled direct comparison of feature-specific cortical representations measured under controlled laboratory conditions and during natural behavior using the same whole-head imaging and analysis pipeline throughout.

Recent perspectives have emphasized the need to move beyond highly controlled laboratory paradigms and study brain function during natural behavior^1,4,55^, and wearable fNIRS has been highlighted as a key technology for this transition^3,29,56,57^. Only a few studies have directly compared neural responses to the same or analogous tasks performed under controlled laboratory and real-world conditions, including studies using mobile EEG^33,34^, and a wearable fNIRS^35^ study comparing prefrontal responses during an n-back task across indoor and outdoor environments. Our study extends this line of work from localized task-related responses and spatially restricted fNIRS measurements to anatomically resolved, whole-head comparisons of feature-specific cortical representations across laboratory and real-world environments.

This comparison relied on two methodological advances. First, cortical responses were analyzed in reconstructed parcel space rather than channel space, enabling anatomically corresponding regions to be compared across participants, conditions, and environments while reducing the dependence of spatial comparisons on channel-level correspondence and optode placement. Second, analogous audiovisual features were extracted from laboratory movies and participant-specific outdoor scene-camera recordings and analyzed within a common modeling framework, allowing representations to be compared across environments despite differences in the sensory measurements available in each setting. Together, these advances provide a practical framework for characterizing cortical representations during natural behavior and for quantifying how feature-related cortical patterns correspond and differ between controlled and real-world environments.

An important consideration in interpreting the present findings concerns the comparison between laboratory movie viewing and outdoor walking. Because these two conditions differ substantially in embodiment, gaze behavior, and locomotor demands, this comparison could be interpreted as contrasting passive and active behavior. However, this was not the aim of the present study. Rather, our objective was to characterize feature-specific cortical organization during natural outdoor behavior and examine how it relates spatially to that measured during laboratory movie viewing. This framing does not eliminate the differences between the two conditions but treats them as an inherent and informative feature of the real-world condition rather than a confound to be equated away^58^. Future work incorporating concurrent physiological and behavioral measurements, including detailed eye-movement metrics, respiration, heart rate, and gait dynamics during outdoor recordings, could help disentangle the extent to which gaze behavior, locomotion, and autonomic state contribute to feature-specific cortical responses, further improving the interpretability of naturalistic feature-brain mapping.

Outdoor luminance estimation represents an additional methodological limitation. Because the eye-tracker scene camera applies automatic exposure correction, luminance could not be recovered from the video recordings; it was instead estimated from the fNIRS dark signal. Indoor and outdoor luminance were therefore derived by methodologically independent means, making it unlikely that their spatial correspondence was driven by a shared luminance-estimation artifact. Nevertheless, future studies using dedicated light sensors could validate and refine the dark-signal approach.

More generally, fNIRS remains vulnerable to systemic physiological and motion-related influences despite channel quality control, TDDR correction, and filtering. Future studies incorporating more explicit modeling of short-separation signals together with concurrent systemic physiological measurements could further improve the physiological specificity of cortical estimates during natural behavior.

The feature space included in the primary mapping analysis was intentionally limited. To preserve interpretability, we intentionally focused on a small set of well-established sensory features and excluded highly correlated predictors, allowing individual cortical response maps to be interpreted more directly. Although this hypothesis-driven approach provided a clear framework for comparing feature-specific cortical organization across laboratory and real-world environments, natural behavior is influenced by a much richer set of correlated visual, auditory, semantic, and behavioral variables. Future studies could extend the present framework to richer feature spaces using approaches such as banded ridge regression, allowing correlated feature groups to be modeled simultaneously while estimating their relative contributions to cortical activity.

Finally, the exploratory encoding analysis had additional limitations. Although the prediction model incorporated five environmental features—optical flow, sound intensity, luminance, body intensity, and face intensity—it did not represent objects present in the scene, actions, social interactions, scene category, or higher-level semantic content. Incorporating frame-level object annotations, action descriptors, captions, or embedding-based representations may capture additional variance, although richer feature spaces will require correspondingly careful regularization and independent evaluation. The models were also fitted separately for each participant; whether an encoding model can generalize across individuals remains an open and more demanding question.

The present study demonstrates that feature-specific cortical representations can be characterized during unconstrained natural real-world behavior using wearable whole-head high-density fNIRS. Comparing cortical response maps derived from controlled movie viewing and unconstrained outdoor walking, we found that representations associated with low-level sensory features — luminance and sound intensity — shared a significant spatial component across environments whereas face-related representations showed no significant cross-environment correspondence, instead engaging distinct cortical patterns in each setting. Feature-specific cortical maps were also reproducible within individuals across repeated outdoor walking sessions, indicating that stable cortical organization can be characterized despite the inherent variability of natural behavior. An exploratory encoding analysis further showed that continuously measured environmental features supported out-of-sample prediction of cortical dynamics during held-out periods of outdoor walking.

Beyond these findings, the combination of wearable whole-head HD-fNIRS, mobile eye tracking, image reconstruction, and parcel-based cortical analysis provides a framework for investigating distributed cortical representations under ecologically valid conditions while maintaining anatomically interpretable comparisons across environments. More broadly, by reconstructing continuous sensory variables from participant-specific recordings, this framework provides a route for relating cortical activity to naturally unfolding experience when the relevant sensory events are not predefined by the experimenter.

## Methods

The study comprised two datasets: (i) naturalistic movie viewing in a controlled laboratory environment (Dataset 1, n = 18) and (ii) outdoor walking (Dataset 2, n = 20). Across both datasets, participants were healthy adults with normal or corrected-to-normal vision and hearing, and the ability to walk independently. All participants were right-handed native English speakers. Exclusion criteria included any current or prior history of neurological trauma, neurological disorders, or psychiatric conditions. All participants provided written informed consent prior to participation. The study was approved by the Institutional Review Board of Boston University.

## Participants

### Dataset 1: Indoor movie-viewing

A total of 22 participants were initially recruited for indoor movie viewing. Three participants were excluded due to technical issues during data acquisition (cable-related signal loss), and one additional participant was excluded due to poor data quality (47.6% of channels failed the quality criteria: 241 for low signal amplitude, 19 saturated, 8 low SNR, 2 outside the source–detector distance range), resulting in a final sample of 18 participants (10 males, 8 females; age range 21–41 years, mean ± SD = 27.1 ± 5.4). Participants viewed a 34-minute segment of a feature film while fNIRS data were recorded.

### Dataset 2: Outdoor walking

Data were initially obtained from 22 subjects. Two participants were excluded from the final analysis due to technical issues with mobile eye-tracking, resulting in a final sample of 20 participants (12 males, 8 females; age range 19–41 years, mean ± SD = 26.7 ± 4.9). Nine of these individuals had also participated in the indoor movie-viewing. Participants in this group completed a multi-stage experimental protocol using simultaneous fNIRS and mobile eye-tracking within a dynamic environment.

## Experimental Paradigm

### Indoor movie-viewing

Participants were seated comfortably in a quiet, dimly lit room throughout the session. Prior to movie presentation, a 6-minute resting-state recording was acquired while participants fixated on a central cross displayed on the screen and were instructed to remain still and relaxed. Following the resting period, participants viewed a 33-minute and 48-second segment (0:00–33:48) of the first movie in the *Back to the Future* series^59^ and were instructed to watch the movie naturally without performing any explicit task. Continuous fNIRS data were recorded throughout the movie presentation. This condition is hereafter referred to as Movie.

The movie was presented using PsychoPy software^60^ (v2022.1.1) on a liquid-crystal display (1,920 × 1,080). Participants were positioned approximately 70 cm from the screen. A trigger signal was sent to the fNIRS acquisition system at movie onset to ensure temporal synchronization between stimulus presentation and neural recordings.

### Outdoor Walking

The outdoor paradigm was conducted within a dynamic and busy campus environment to capture neural responses during real-world behavior. All sessions were conducted during daylight hours between 10:00 and 16:00.

Prior to the first walking run, participants remained standing for 2 min. They then walked at their own pace along a predefined route (Walking 1), remained seated for 10 min at the turnaround point (Sitting; used only in the prediction analysis), and returned along the same path (Walking 2). The route covered approximately 0.64 km in each direction (1.3 km total) through the campus area (Fig. 1d). The route was introduced to the participants immediately before the recording to ensure they could navigate independently while maintaining natural behavior. Each walking run lasted approximately 10 min, depending on individual walking speed. The environment was highly dynamic, with pedestrian and vehicular traffic and multiple street crossings. Participants were instructed to walk naturally and not to speak during the recordings.

Continuous fNIRS and mobile eye-tracking data were acquired throughout all outdoor phases. The experimenter followed the participant at a distance to ensure data integrity while minimizing interference.

## Data Acquisition

### fNIRS system and probe configuration

For indoor recordings, fNIRS data were collected using a whole-head high-density probe layout (Fig. 1b,c) developed for the NinjaNIRS2022 system, comprising 56 sources and 144 detectors and yielding 567 measurement channels^61^. Data were acquired at two wavelengths (760 nm and 850 nm), resulting in a total of 1,134 measurements. The cap was positioned using the international 10-20 system as reference. The fNIRS sampling rate was 8.988 Hz.

For outdoor recordings, an updated 2024 version of the NinjaNIRS system was used. This version operated under the same acquisition principles but was configured as a backpack-based, wireless unit to allow natural, unconstrained movement. Both the acquisition computer and the fNIRS units were connected through a common Wi-Fi hotspot, and outdoor data were acquired using the NinjaWeb interface. To reduce ambient light interference, a light-blocking cap was placed over the fNIRS cap, with dark signal levels monitored during system setup.

### Eye Tracking

During outdoor sessions, mobile eye-tracking was performed using the Pupil Labs Neon system (Pupil Labs GmbH, Berlin, Germany). The integrated scene camera recorded the participant’s point-of-view (POV) video at 1600 × 1200 pixels and 30 Hz with a 132° × 81° field of view, while environmental audio was captured simultaneously via the device’s built-in microphones. The POV video stream provided a participant-specific record of the audiovisual environment and was processed for feature extraction using the same framework applied to the indoor movie. The system operates without the need for manual calibration. The device connects to a companion mobile device that enables recording and data storage.

fNIRS and eye-tracking recordings were temporally synchronized via cross-correlation of acceleration signals from two inertial measurement units (IMUs): one embedded in the eye tracker and a separate head-mounted IMU used with the fNIRS system. A head movement performed at the start of each run provided an additional alignment marker.

### Stimulus Feature Extraction

Time-varying visual and auditory features were extracted from the indoor movie stimulus and the outdoor eye-tracking video recordings using automated computer vision and audio analysis methods. Features were selected to span both low-level (luminance, sound intensity, optical flow) and higher-level (face intensity, body intensity, speech) sensory features widely used in feature-based naturalistic neuroimaging^20,22,62^.

### Auditory features

Sound intensity was estimated using a multiband log-power envelope approach. The audio signal was decomposed using a short-time Fourier transform (STFT; FFT length = 2048; hop size = 50 ms). Spectral power was computed within contiguous 200-Hz frequency bands spanning 200–5000 Hz^22^. Band-specific power values were averaged across all frequency bands to obtain a broadband power envelope, which was subsequently converted to decibel (dB) units to yield the final sound intensity time series. The same procedure was applied to both datasets.

Human speech presence was detected in the indoor movie audio using the pre-trained Silero voice activity detection (VAD) model (v5)^63^. The model output was thresholded at 0.5 to classify audio frames as speech or non-speech; speech frames were coded as 1 and non-speech frames as 0, yielding a binary time series representing speech presence over time.

### Visual features

Luminance and optical flow were extracted as low-level features, and face and body metrics as higher-level features. For the indoor dataset, luminance was computed frame-wise from sRGB video after conversion to linear RGB, according to the ITU-R BT.709 definition^64^. The luminance regressor was obtained by averaging relative luminance across all pixels in each frame. For the outdoor dataset, direct estimation of luminance from the eye-tracking videos was not feasible because of automatic exposure adjustment^65^. Luminance was therefore indexed by the fNIRS detector dark signal rather than by the video itself. Averaging the dark signal across detectors yielded the outdoor luminance regressor, which tracked changes in ambient illumination along the walking route.

Optical flow was estimated for both datasets using the Farnebäck dense optical flow algorithm^66^, with motion magnitude averaged across pixels to obtain a global motion time series. For the outdoor videos, camera motion induced by head and body movement was estimated and removed prior to flow computation.

Face and body detection were performed using the YuNet face detector^67^ and the pre-trained YOLOv8 object detector^68^ (trained on the COCO dataset^69^), respectively, applied to every frame. For both the indoor and outdoor datasets, face intensity was computed as the total area of all detected face bounding boxes divided by the total frame area, following studies that quantify face content continuously^39^. This measure incorporates all detected faces within the scene rather than a single dominant face, capturing the cumulative visual presence of multiple individuals — common in outdoor environments where no single face is emphasized. Similarly, body intensity was computed as the total area of all detected person bounding boxes divided by the total frame area.

All extracted features were resampled to match the fNIRS acquisition rate and temporally aligned with the neural recordings. For the indoor dataset, sound intensity was floored at its 1st percentile and face intensity capped at its 99th percentile to remove outliers; no clipping was applied outdoors.

### fNIRS preprocessing

All fNIRS processing was performed using Cedalion toolbox^70^ (v25.1.1). Raw light intensity signals were converted to optical density (OD). Channel quality was assessed using amplitude, source–detector distance, and signal-to-noise ratio (SNR) criteria. Channels were flagged as low-quality if they were saturated (mean intensity above 0.84 V), exhibited a mean intensity below 1 × 10⁻³ V in either wavelength or fell outside the source–detector distance range of 1–40 mm. Low-quality channels were down-weighted in subsequent reconstruction by inflating their measurement variance, thereby preserving consistent spatial indexing across subjects. Motion artifacts were corrected using Temporal Derivative Distribution Repair (TDDR)^71^. OD signals were then bandpass filtered (0.01–0.5 Hz) to reduce slow drifts and high-frequency physiological noise.

### Image reconstruction

Image reconstruction was performed using a diffuse optical tomography (DOT) framework based on an indirect reconstruction formulation, in which OD changes were reconstructed separately for each wavelength and subsequently converted to chromophore concentrations. Reconstruction was performed without spatial basis expansion, as indirect reconstruction without spatial basis functions has been shown to reduce oxy–deoxy crosstalk and improve spatial localization accuracy in high-density hexagonal measurement arrays^72^.

For each wavelength *λ*, the measurement model was defined as:

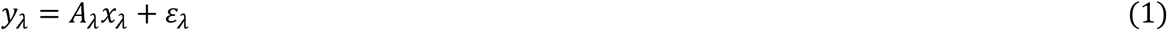

where *y_λ_* denotes the channel-wise OD measurements, *A_λ_* is the sensitivity matrix (forward model) and *x_λ_* represents the unknown vertex-wise OD changes in image space. The sensitivity matrix A was generated from Monte Carlo photon propagation model from pmcx^73^ (10⁸ photons per optode) within a two-surface head model derived from the ICBM152 anatomical template^74^. The cortical surface contained 15,002 vertices, which served as reconstruction nodes in image space, while the scalp surface contained 10,018 vertices.

Spatial regularization was defined based on the column energy of the forward model

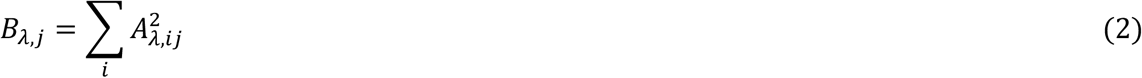

where *i* indexes measurement channels and *j* indexes cortical vertices. A spatial prior was constructed as

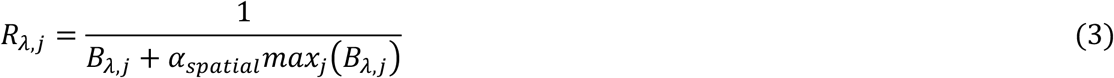

Where *α_spatial_* controls the degree of the spatial regularization and *max*_j_5*B_λ_*_,j_) denotes the maximum column energy across cortical vertices at wavelength *λ*.

The regularized reconstruction operator was computed as:

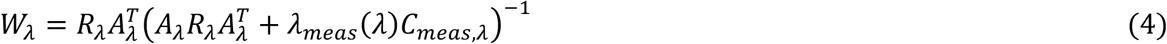

where *C_meas_*_,*λ*_ is a diagonal measurement covariance matrix derived from the variance of the temporal derivative of the optical density signals for each channel and wavelength, allowing measurements with higher noise levels to be down-weighted during inversion. The spatial regularization parameter was set to α_spatial = 10⁻³, the value that maximizes the area under the normalized singular value spectrum when no spatial basis functions are used^72^. The measurement regularization term, *λ*_meas_ = *α*_meas_*λ_R_λ*_max_, was set to approximately one, with *α*_meas_ = 10^4^ and *λ_R_* = 10^−6^ following the optimization reported for this probe geometry^72^. The reconstructed optical density image at each wavelength was obtained as

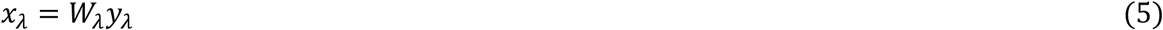

The wavelength specific OD images were subsequently converted to chromophore concentration changes using the inverse extinction coefficient matrix,

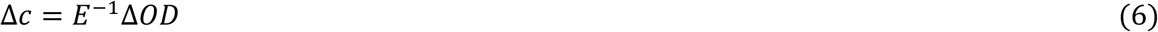

where Δ*OD* = [*x_λ_*_1_, *x_λ_*_2_]^T^, Δ*c* = [Δ*HbO*, Δ*HbR*]*^T^* and *E* is the extinction coefficient matrix relating wavelength-specific optical density changes to HbO and HbR concentration changes.

This reconstruction yielded vertex-level estimates of HbO and HbR concentration changes across the cortical surface, which were used for subsequent analyses.

Following image reconstruction, statistical modelling was performed in parcel space using the reconstructed HbO and HbR time series. Vertex-level reconstructed time series were averaged within cortical parcels defined by the Schaefer-600 parcellation^75^, using the parcel labels of the ICBM152 forward model, yielding parcel-level time series (time × parcel × chromophore) across 602 parcels for each subject, two of which correspond to the medial walls.

## Feature-based GLM analysis

### HRF modelling

Each feature time series was padded by 30 s to minimize edge effects and convolved with a single-gamma hemodynamic response function (τ = 0 s, σ = 3.0 s) to account for the delayed hemodynamic response, then normalized to the [0,1] range using min–max scaling prior to statistical modelling (Fig. 2b).

### Regressor collinearity and feature selection

Pairwise correlations were computed among the HRF-convolved and normalized feature regressors to assess multicollinearity and guide feature selection. Regressor pairs with |*r*| > 0.5 were considered to exhibit substantial collinearity and were evaluated for feature exclusion. Body intensity exhibited consistently high correlations with the face intensity regressor in both the indoor and outdoor datasets (Fig. 2c), whereas optical flow showed high correlations with the luminance regressor in a subset of outdoor participants (Supplementary Fig. 1). To reduce collinearity and improve the interpretability of the regression coefficients, the body intensity and optical flow regressors were excluded from the GLM analyses. Face intensity and luminance were retained because cortical responses to faces and low-level visual features are well characterized in the naturalistic neuroimaging literature, providing an established basis for the a priori regions of interest used here^12,76^. The final design matrix for the Movie condition therefore included luminance, sound intensity, speech presence, and face intensity, whereas the Walking models included luminance, sound intensity, and face intensity. The speech regressor was not included in the Walking models because speech encountered during free walking was sparse and incidental and was not directly comparable to the structured dialogue in the indoor movie. Feature-mapping analyses were restricted to Walking 1 and Walking 2, where participants actively moved through the environment and encountered continuously changing sensory input; the Sitting period was excluded because environmental feature variation was substantially reduced while participants remained stationary.

### General Linear Model

A general linear model (GLM) was used to estimate the relationship between feature regressors and the parcel-level hemodynamic response (Fig. 8c). The model was specified as:

**Fig. 8.**
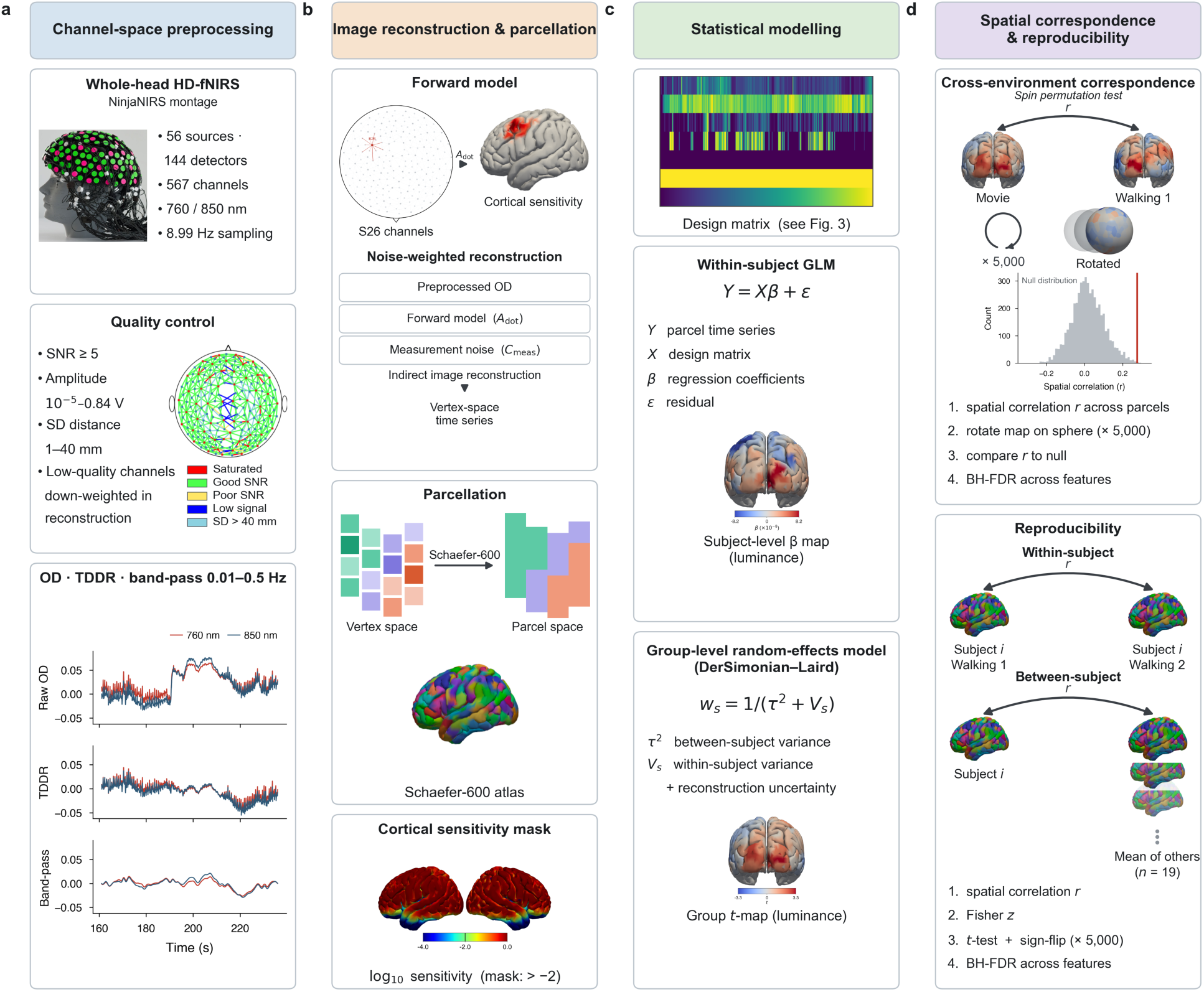
Overview of the whole-head HD-fNIRS processing and analysis pipeline. **a,** Channel-space preprocessing comprised channel quality assessment followed by conversion to optical density, motion correction (TDDR), and band-pass filtering (0.01–0.5 Hz). Channels failing quality criteria were down-weighted rather than discarded, entering image reconstruction with inflated measurement noise. **b,** Preprocessed optical density was projected from channel space to cortical vertex space by indirect image reconstruction, weighted by the measurement noise covariance *C*meas. A representative source (S26) is highlighted to illustrate the channel-to-cortex mapping. Vertex-wise time series were averaged within Schaefer-600 parcels, and analyses were restricted to parcels with log₁₀ of the channel-summed sensitivity at 850 nm above −2. **c,** Feature-specific responses were estimated with parcel-wise within-subject GLMs, yielding subject-level β maps (shown for a representative participant). Group estimates used a DerSimonian–Laird random-effects model in which within-subject variance was inflated by the reconstruction posterior variance, yielding group *t*-maps per feature. **d,** Cross-environment correspondence between movie and outdoor walking maps was assessed by spin permutation (× 5,000), which preserves cortical spatial autocorrelation. Within-subject reproducibility (two walking runs) and between-subject reproducibility (each participant against the mean of the remaining participants) were quantified as spatial correlations across parcels, Fisher *z*-transformed and tested against zero using sign-flip permutation tests. Multiple comparisons across features were controlled by Benjamini–Hochberg FDR.

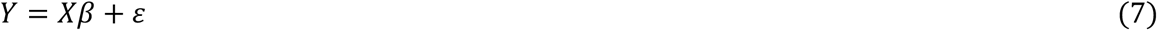

where Y is the parcel-level HbO time series (time × parcel), X is the design matrix containing the HRF-convolved feature regressors, a first-order polynomial drift term, and a weighted global mean regressor, β represents the feature-specific regression coefficients, and ε is the residual error. The weighted global mean regressor was computed as a weighted average across cortical parcels, with parcel weights derived from image reconstruction sensitivity and measurement noise. The model was fitted independently to each cortical parcel using ordinary least squares. Feature-specific β coefficients and their associated covariance matrix (Σβ) were estimated independently for each cortical parcel and retained for subsequent group-level analyses.

## Group Level Analysis

Group-level inference was performed using a DerSimonian–Laird random-effects meta-analysis^77^. Subject-specific feature regression coefficients were combined across participants using inverse-variance weighting. Subject weights were computed as

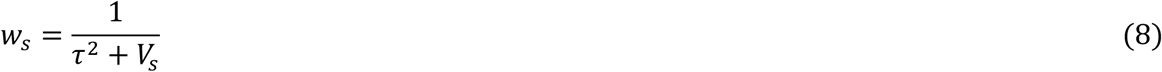

where *V_s_*is the within-subject variance estimate derived from the GLM covariance matrix and *τ*^2^is the between-subject variance estimated using the DerSimonian–Laird method. The group-level regression coefficient was then calculated as

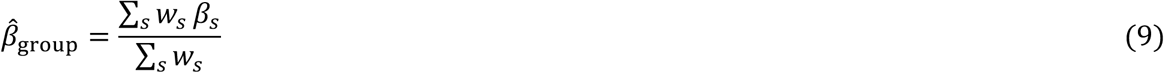

where *β_s_*is the subject-specific regression coefficient. Within-subject variance estimates were obtained from the GLM covariance matrix and scaled by a factor reflecting the ratio of total to residual variance, so that the additional uncertainty introduced by the image reconstruction was propagated into the group-level weighting. Group-level standard errors and corresponding t statistics were computed independently for each cortical parcel and feature regressor, yielding feature-specific cortical t maps for subsequent analyses.

## Feature-specific spatial correlation analysis

Feature-specific group-level cortical *t* maps obtained from each experimental condition (Movie, Walking 1, Walking 2) were compared to assess the spatial correspondence of cortical response patterns. Spatial similarity between corresponding *t* maps was quantified using parcel-wise Pearson correlation coefficients. Whole-brain analyses were restricted to cortical parcels with sufficient image reconstruction sensitivity (*log*(∑*_c_ A_c_* > −2) at 850 nm; Fig. 8b). Because cortical maps exhibit substantial spatial autocorrelation, statistical significance was assessed using a spin permutation test^78^ rather than conventional parametric significance testing (Fig. 8d). During each permutation, cortical parcel coordinates were projected onto the fsaverage sphere^79^ and randomly rotated while preserving hemispheric topology. The left hemisphere was randomly rotated, and the corresponding right-hemisphere rotation was generated using a mirrored transformation to preserve contralateral symmetry. Parcel values were then reassigned using nearest-neighbor matching, and a null distribution of spatial correlation coefficients was generated from 5,000 rotations. Two-sided empirical *P* values were calculated as the proportion of permuted correlations whose absolute value exceeded the observed correlation. Resulting *P* values were corrected for multiple comparisons using the Benjamini–Hochberg false discovery rate procedure^80^ (*q* = 0.05), applied separately to the six movie–walking comparisons and the three walking–walking comparisons.

### ROI-based spatial correlation analysis

To evaluate the spatial correspondence of feature-specific cortical response patterns within functionally relevant regions, ROI-based analyses were additionally performed. ROIs were defined a priori based on previous neuroimaging literature.

The luminance ROI encompassed striate and early extrastriate visual cortex (BA17, BA18, and BA19), reflecting evidence that low-level visual features engage early visual areas beyond primary cortex during naturalistic viewing^36,37^. The sound intensity ROI encompassed primary and temporal association auditory cortex (BA41, BA42, BA22, and BA21), spanning the hierarchical cortical organization involved in processing continuous acoustic features^38^. The face intensity ROI comprised cortical regions consistently associated with face perception, including the fusiform face area (BA37)^76^ and the posterior superior temporal sulcus (pSTS)^81,82^.

Schaefer parcels were assigned to Brodmann areas according to their dominant anatomical label. Spatial correlations between corresponding group-level t maps were computed within each ROI, and statistical significance was assessed using the spin permutation framework and BH-FDR correction described above.

## Reproducibility analyses

Subject-level spatial correlations were performed to evaluate the reproducibility of feature-specific cortical response patterns at the individual participant level. Correlations were computed between feature-specific *t* maps across the whole-brain sensitivity-masked parcel set and included both within-subject and between-subject analyses.

### Within-subject spatial correlations

For the outdoor walking experiment, each participant completed two walking runs along the same route. Feature-specific subject-level *t* maps were estimated independently for each run, and parcel-wise Pearson correlations were computed between the corresponding maps.

For the Movie experiment, direct test–retest comparisons were not possible because each participant viewed the movie only once. Therefore, the ∼34-min movie was divided into two non-overlapping halves. Separate GLMs were fitted to each half using the corresponding feature regressors. The HRF convolution was applied to the uninterrupted time series, and model estimation was restricted to the time points belonging to each half. Subject-level *t* maps were then estimated independently for each half, and parcel-wise Pearson correlations were computed between the corresponding maps (Fig. 8d).

Individual correlation coefficients were transformed using Fisher’s r-to-z transformation, and statistical significance was assessed using a two-sided sign-flip permutation test (5,000 permutations). Resulting *P* values were corrected across features within each condition using the Benjamini–Hochberg false discovery rate procedure (*q* = 0.05).

### Between-subject spatial correlations

Between-subject reproducibility was assessed using LOO approach based on feature-specific subject-level *t* maps. The LOO approach avoids the statistical dependency inherent in pairwise comparisons by comparing each participant with a group map estimated from the remaining participants^51^.

For the Movie experiment, each participant’s feature-specific *t* map was spatially correlated with the corresponding LOO group *t* map estimated from the remaining 17 participants. For the outdoor walking experiment, the two runs were analyzed independently; within each run, each participant’s t map was correlated with the LOO group t map estimated from the remaining 19 participants (Fig. 8d).

Correlation coefficients were transformed using Fisher’s *r*-to-*z* transformation. Statistical significance was assessed using a two-sided sign-flip permutation test (5,000 permutations) performed on the Fisher-transformed correlation coefficients. Resulting *P* values were corrected across features within each condition using the Benjamini–Hochberg false discovery rate (BH-FDR) procedure (*q* = 0.05).

### Naturalistic outdoor fNIRS encoding model

An exploratory subject-specific encoding analysis was performed using ridge regression to predict parcel-level HbO responses from time-lagged environmental features. Walking 1, Sitting, and Walking 2 were included in model development to use the available continuous outdoor recordings, whereas out-of-sample evaluation was restricted to the two walking runs, consistent with the primary focus on prediction during dynamic real-world behavior. In each outer fold, the held-out walking half was reserved exclusively for testing, while the remaining eligible samples constituted the development set used for model selection and fitting. Analyses were performed independently within participant; no brain data were pooled across participants for model fitting.

### fNIRS preprocessing for prediction analysis

fNIRS preprocessing followed the same general framework described above, including optical-density conversion, channel-quality assessment, TDDR motion correction, and temporal filtering, with analysis-specific modifications to channel-quality criteria and filtering parameters as detailed below.

Continuous-wave intensity data were read from SNIRF files and restricted to channels represented in the subject-independent sensitivity matrix. Nonfinite or nonpositive intensity samples were replaced by a small positive value before quality assessment and optical-density conversion. Channel quality was evaluated independently in every run using both wavelengths. A channel was retained only when (i) its mean raw intensity was between 10^−3^ and 0.84 V, (ii) its signal-to-noise ratio, defined as the temporal mean divided by the temporal standard deviation, was finite and exceeded 5, and (iii) its raw intensity had a nonzero temporal range. Both wavelengths were required to satisfy every criterion. Retained intensity measurements were converted to optical density. Motion correction was applied independently to each channel and wavelength using TDDR^71^. Optical-density time courses were then zero-phase Butterworth filtered from 0.005 to 0.5 Hz.

### Image reconstruction and parcel time courses for prediction analysis

Optical-density measurements were reconstructed on the ICBM152 cortical head model with the same setting as described in Image reconstruction section. Sensitivity was evaluated at every cortical vertex separately for both wavelengths. A vertex was considered measurable when the log10 summed channel sensitivity exceeded −2 at both wavelengths and the vertex belonged to the cortical brain mask. Reconstructed vertex time courses were averaged within atlas parcels. A parcel was retained when it contained at least 10 sensitive vertices, at least 25% of its cortical vertices were sensitive, and it satisfied these requirements in all three runs for that participant. Only retained parcels entered prediction and statistical summaries.

### Naturalistic stimulus features

The encoding analysis used five synchronized features: (1) optical flow, (2) sound intensity, (3) luminance, (4) body intensity, and (5) face intensity. Unlike the feature-specific GLM analysis, all five features were retained for prediction because the objective was out-of-sample prediction rather than interpretation of individual regression coefficients, with correlated predictors handled through ridge regularization. The original feature streams were sampled at approximately 8.99 Hz. Before downsampling, every feature was filtered with a fourth-order, zero-phase 0.5-Hz low-pass Butterworth filter to prevent aliasing and to match the upper frequency bound of the processed HbO response. Filtered features and parcel HbO were then interpolated independently onto their common SNIRF interval at 2 Hz. Resampling and temporal modeling were performed separately within every run, preventing FIR predictors from crossing run boundaries.

### FIR encoding model

For feature *k*, the primary finite-impulse-response design contained 12 delayed copies spanning 1–12 s:

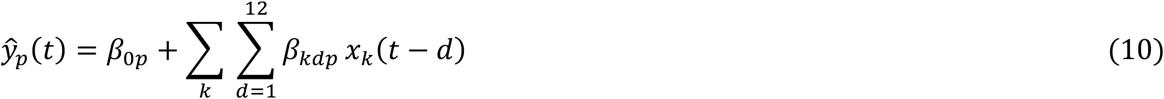

where *ŷ_p_*(*t*) is predicted HbO in parcel *p*, *x_k_* is stimulus feature *k*, and *β_kdp_* is its parcel-and delay-specific coefficient^8,83^. Five features produced 60 predictors. The initial 32 s of every run were excluded after temporal-design construction so that all temporal-model comparisons used an identical initial trim.

Ordinary multi-output ridge regression was fitted separately within participant^83^. Model selection was nested inside each outer test fold. Specifically, the outer-fold development set was divided into five contiguous blocked cross-validation folds^84^. In each inner cross-validation iteration, four blocks were used for coefficient estimation and the remaining block was used for validation; the validation block rotated so that every development block served once as validation. A 12-s gap was removed around every validation block. Predictor and response standardization parameters were estimated from the inner training blocks and applied to the corresponding validation block, preventing validation information from influencing scaling.

The final outer-fold model was then evaluated once on the untouched held-out walking half. Thus, inner validation data were used only for hyperparameter selection, whereas the outer held-out half was used only to estimate generalization performance.

### Four-fold nested blocked evaluation

Each walking run was divided at its midpoint into nonoverlapping early and late halves, producing four outer test folds: walking 1 early, walking 1 late, walking 2 early, and walking 2 late. Usable held-out durations ranged from 207.5 to 360 s after synchronization and FIR trimming. For a given outer fold, the selected walking half was withheld completely from model development.

More precisely, the sitting run, other walking run, and nonheld-out half of the selected walking run constituted the outer-fold development set, rather than a single undivided training set. Model selection and final testing followed a nested structure: the five inner folds select the ridge penalty, while the four outer walking-half folds measure held-out prediction accuracy^84^. The outer test half never contributed to feature or response standardization, ridge-penalty selection, or coefficient estimation. A 30-s buffer adjoining the outer held-out boundary was also removed from the development set. Every participant completed all four outer folds.

Prediction accuracy was Pearson correlation between predicted and observed HbO, calculated independently for every retained parcel and fold. Parcel correlations were clipped only for numerical stability and transformed using Fisher’s *z* = arctanh(*r*). Within each participant, Fisher-z values were averaged across parcels and the four folds and transformed back to correlation units. The primary cohort statistic was the arithmetic mean of these 20 participant-level correlations.

### Confidence interval and temporal-shift null

A nonparametric 95% confidence interval was obtained by resampling participants with replacement 100,000 times and recalculating the cohort mean. Chance performance under preserved temporal autocorrelation was assessed by circularly shifting each held-out prediction relative to its observed HbO. Shifts with circular distance below 60 s were excluded. For each of 10,000 iterations, one permissible shift was independently drawn for every participant and fold, parcel correlations were Fisher averaged using the same hierarchy as the primary statistic, and the cohort mean was recalculated. The one-sided Monte Carlo p-value used the plus-one correction:

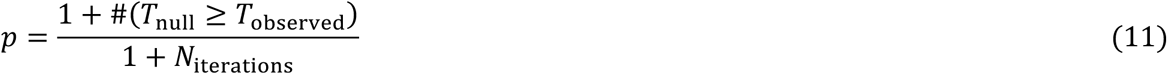

### Correct-segment specificity control

To test whether predictions matched their corresponding experience rather than generic slow hemodynamics, each held-out prediction was additionally paired with equal-length HbO windows from the same participant’s other real walking data. Sitting data were excluded from this control. Candidate windows advanced in 60-s increments and could not overlap the prediction’s true held-out segment. This yielded 422 unique mismatched prediction–HbO pairings across participants and folds. In each of 10,000 cohort-null iterations, one mismatched window was sampled for every participant and fold, followed by the same Fisher-z aggregation used for the primary statistic. This procedure preserved genuine participant-specific HbO dynamics while disrupting segment identity.

### Spatial maps and multiplicity

For visualization, parcel correlations were Fisher averaged across four folds within participant and then across participants for the group descriptive map. The cohort map was restricted to the 247 parcels satisfying the sensitivity and quality-control criteria in all 20 participants; parcels lacking complete participant coverage were displayed as missing. Network summaries were calculated by Fisher averaging the complete-case parcel correlations within each network separately for every participant, yielding 20 subject-level values per represented network. These spatial and network summaries are descriptive: no parcel or network was labeled statistically significant. The single prespecified cohort-level primary test did not require false-discovery-rate correction. Any future inferential claims about individual parcels or networks will require appropriate multiplicity control.

## Acknowledgements

This project was supported by the National Institutes of Health (Grant No. U01-EB 029856). We are especially grateful to Alexander von Lühmann for the idea of the protective cap for the mobile fNIRS system, and to Yüksel Yücel for skillfully bringing the idea to life by making the cap. We also thank Sudan Duwadi for his assistance during pilot recordings and Sharayu Shrinivas for her assistance with data collection.

