## Supplementary Information for "The Walking Brain: Mapping feature-specific cortical responses to naturally unfolding sensory experience in the real world"

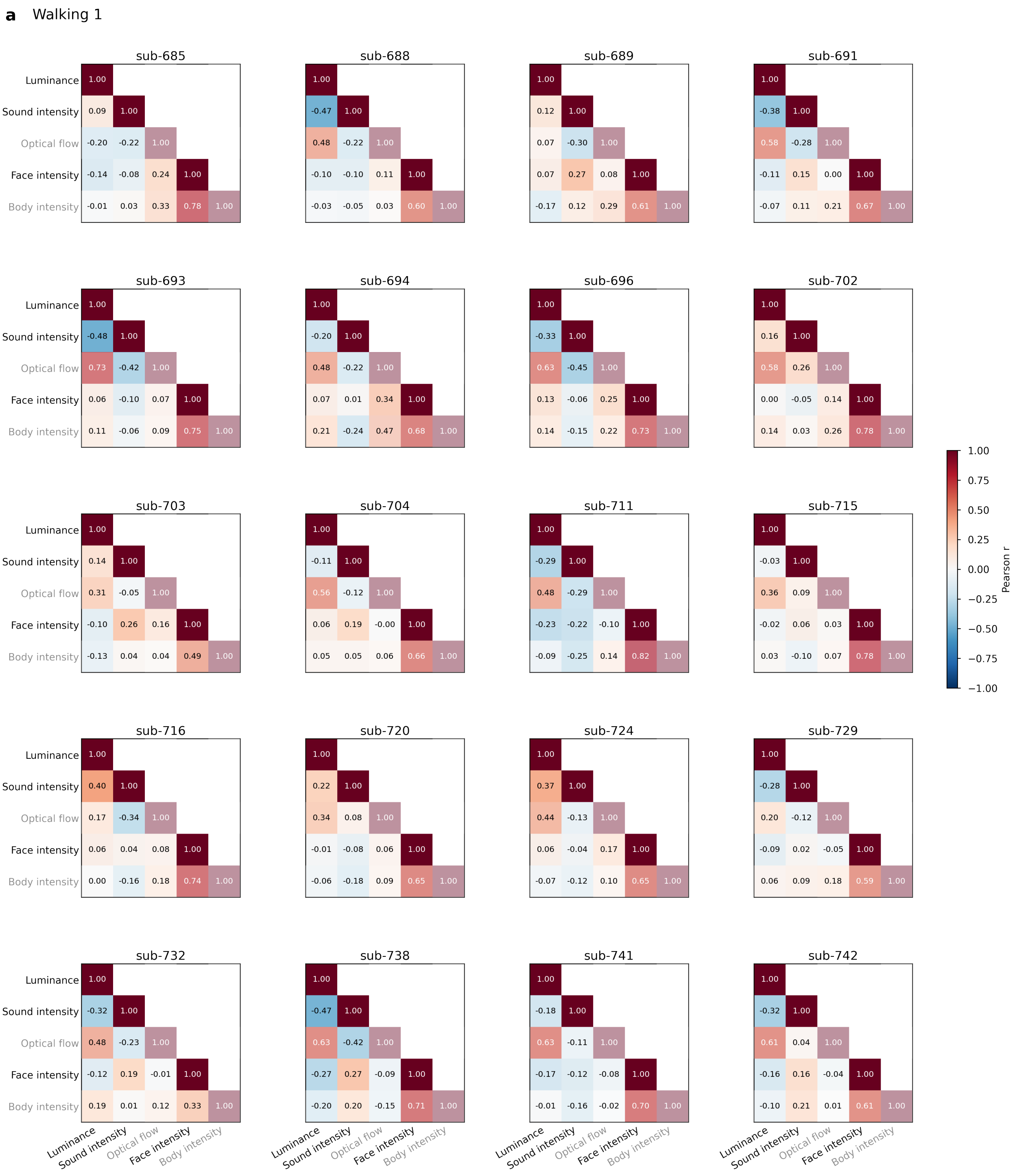

**b** Walking 2

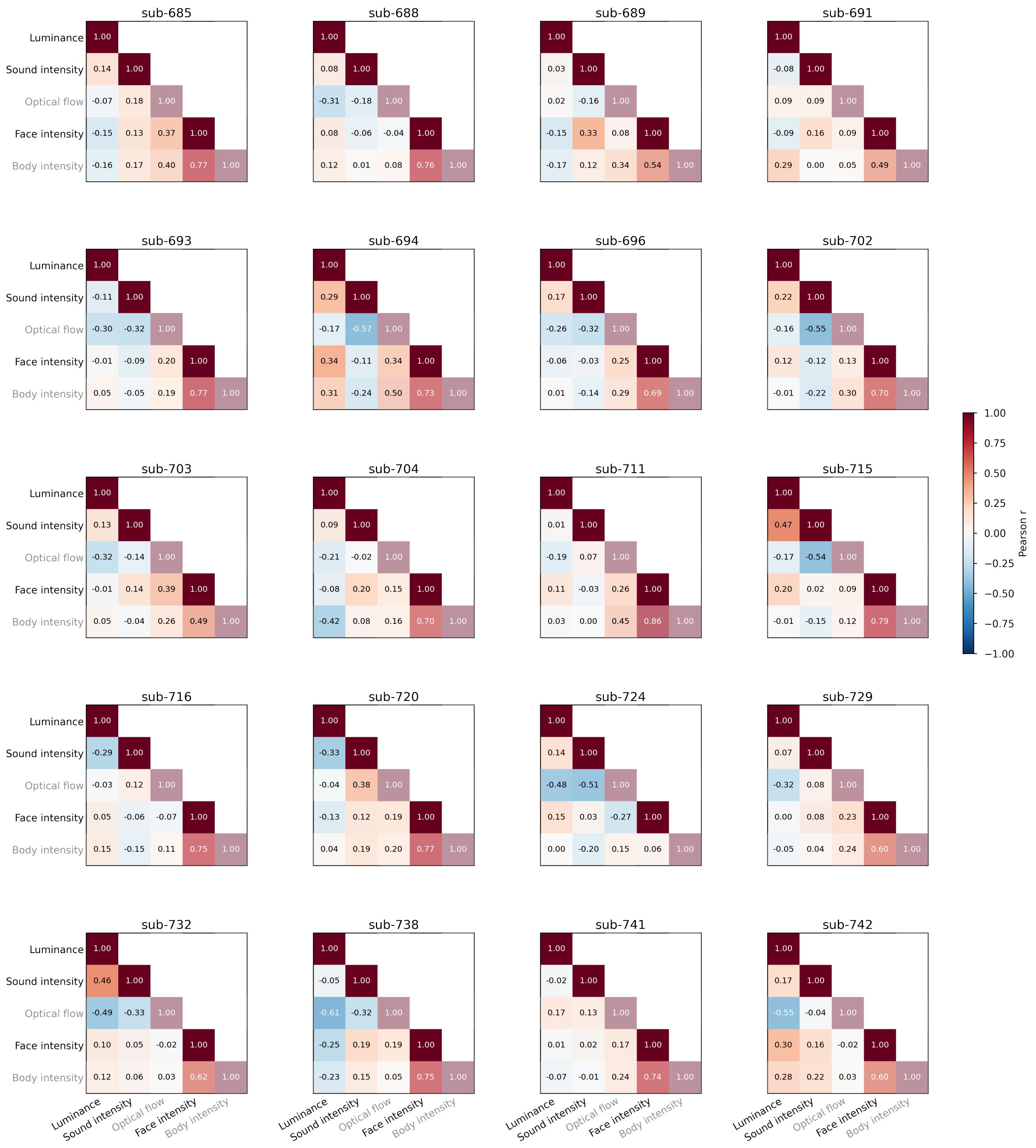

Luminance
Sound intensity
Optical flow
Face intensity
Body intensity
sub-685
1.00
0.14 1.00
-0.07 0.18 1.00
-0.15 0.13 0.37 1.00
-0.16 0.17 0.40 0.77 1.00
sub-688
1.00
0.08 1.00
-0.31 -0.18 1.00
0.08 -0.06 -0.04 1.00
0.12 0.01 0.08 0.76 1.00
sub-689
1.00
0.03 1.00
0.02 -0.16 1.00
-0.15 0.33 0.08 1.00
-0.17 0.12 0.34 0.54 1.00
sub-691
1.00
-0.08 1.00
0.09 0.09 1.00
-0.09 0.16 0.09 1.00
0.29 0.00 0.05 0.49 1.00
sub-693
1.00
-0.11 1.00
-0.30 -0.32 1.00
-0.01 -0.09 0.20 1.00
0.05 -0.05 0.19 0.77 1.00
sub-694
1.00
0.29 1.00
-0.17 -0.57 1.00
0.34 -0.11 0.34 1.00
0.31 -0.24 0.50 0.73 1.00
sub-696
1.00
0.17 1.00
-0.26 -0.32 1.00
-0.06 -0.03 0.25 1.00
0.01 -0.14 0.29 0.69 1.00
sub-702
1.00
0.22 1.00
-0.16 -0.55 1.00
0.12 -0.12 0.13 1.00
-0.01 -0.22 0.30 0.70 1.00
sub-703
1.00
0.13 1.00
-0.32 -0.14 1.00
-0.01 0.14 0.39 1.00
0.05 -0.04 0.26 0.49 1.00
sub-704
1.00
0.09 1.00
-0.21 -0.02 1.00
-0.08 0.20 0.15 1.00
-0.42 0.08 0.16 0.70 1.00
sub-711
1.00
0.01 1.00
-0.19 0.07 1.00
0.11 -0.03 0.26 1.00
0.03 0.00 0.45 0.86 1.00
sub-715
1.00
0.47 1.00
-0.17 -0.54 1.00
0.20 0.02 0.09 1.00
-0.01 -0.15 0.12 0.79 1.00
sub-716
1.00
-0.29 1.00
-0.03 0.12 1.00
0.05 -0.06 -0.07 1.00
0.15 -0.15 0.11 0.75 1.00
sub-720
1.00
-0.33 1.00
-0.04 0.38 1.00
-0.13 0.12 0.19 1.00
0.04 0.19 0.20 0.77 1.00
sub-724
1.00
0.14 1.00
-0.48 -0.51 1.00
0.15 0.03 -0.27 1.00
0.00 -0.20 0.15 0.06 1.00
sub-729
1.00
0.07 1.00
-0.32 0.08 1.00
0.00 0.08 0.23 1.00
-0.05 0.04 0.24 0.60 1.00
sub-732
1.00
0.46 1.00
-0.49 -0.33 1.00
0.10 0.05 -0.02 1.00
0.12 0.06 0.03 0.62 1.00
sub-738
1.00
-0.05 1.00
-0.61 -0.32 1.00
-0.25 0.19 0.19 1.00
-0.23 0.15 0.05 0.75 1.00
sub-741
1.00
-0.02 1.00
0.17 0.13 1.00
0.01 0.02 0.17 1.00
-0.07 -0.01 0.24 0.74 1.00
sub-742
1.00
0.17 1.00
-0.55 -0.04 1.00
0.30 0.16 -0.02 1.00
0.28 0.22 0.03 0.60 1.00
1.00
0.75
0.50
0.25
0.00
−0.25
−0.50
−0.75
−1.00
Pearson r

**Supplementary Fig. 1. | Regressor collinearity in individual participants during outdoor walking.** **a** Pearson correlation matrices between HRF-convolved stimulus regressors for each participant (n = 20) during Walking 1. **b** The same, for Walking 2. Regressors were convolved with the HRF used in the GLM and min–max normalized before correlation. Face and body regressors were consistently correlated across participants and both runs, reflecting the co-occurrence of faces and bodies in the outdoor scene. Luminance and optical flow were also correlated in several participants, reaching r = 0.73. Because of this collinearity, body intensity and optical flow were not retained in the final model, which included luminance, sound intensity and face intensity as separable predictors.

**Supplementary Table 1 | Spatial correlation between feature-specific group t maps.** Spin-permutation *p* values, uncorrected and FDR-adjusted; *adjusted $p < 0.05$. W1, Walking 1; W2, Walking 2.

| Feature | Comparison | r | p | p (FDR) |
|---|---|---|---|---|
| **Whole brain** | | | | |
| Luminance | Movie – W1 | 0.276 | < 0.001 | 0.001* |
| | Movie – W2 | 0.030 | 0.741 | 0.741 |
| | W1 – W2 | 0.285 | < 0.001 | < 0.001* |
| Sound intensity | Movie – W1 | 0.239 | 0.023 | 0.046* |
| | Movie – W2 | 0.321 | 0.001 | 0.003* |
| | W1 – W2 | 0.055 | 0.584 | 0.584 |
| Face intensity | Movie – W1 | 0.111 | 0.166 | 0.250 |
| | Movie – W2 | 0.054 | 0.482 | 0.578 |
| | W1 – W2 | 0.116 | 0.196 | 0.294 |
| **ROI** | | | | |
| Luminance | Movie – W1 | 0.516 | 0.004 | 0.011* |
| | Movie – W2 | 0.497 | 0.006 | 0.011* |
| | W1 – W2 | 0.620 | < 0.001 | < 0.001* |
| Sound intensity | Movie – W1 | 0.354 | 0.115 | 0.173 |
| | Movie – W2 | 0.634 | < 0.001 | < 0.001* |
| | W1 – W2 | 0.380 | 0.022 | 0.032* |
| Face intensity | Movie – W1 | 0.060 | 0.838 | 0.838 |
| | Movie – W2 | 0.235 | 0.331 | 0.398 |
| | W1 – W2 | 0.336 | 0.147 | 0.147 |

**a**

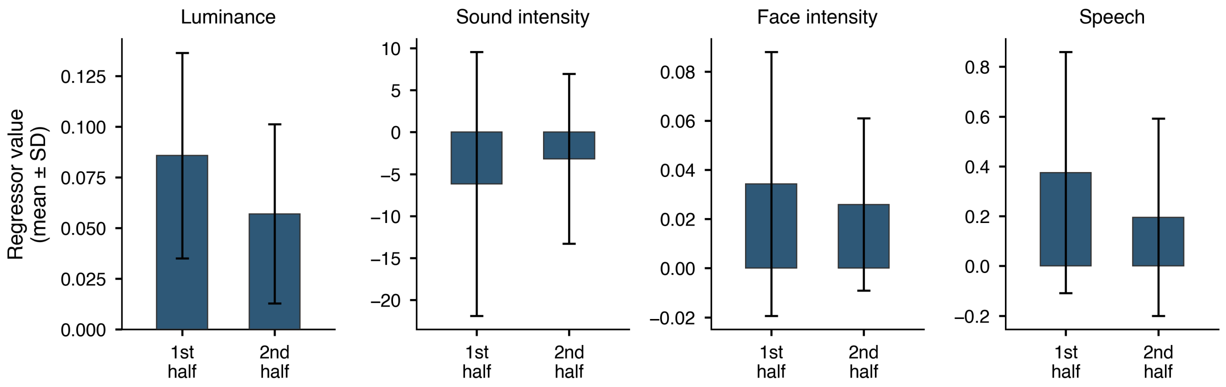


**b**

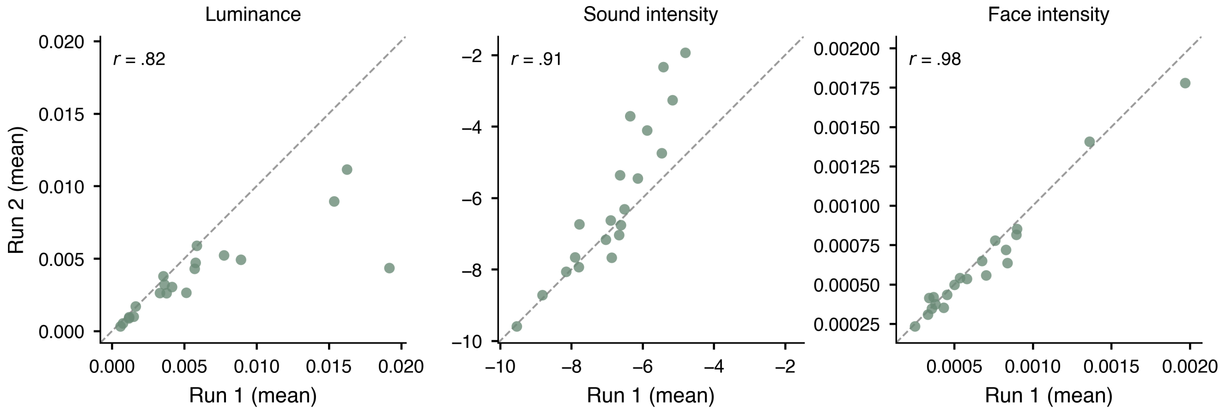


**Supplementary Fig. 2. | Feature content across the two portions of each recording. a,** Movie: mean ± SD of each regressor in the first and second half. All participants viewed the same movie, so a single value pair is shown per feature. **b,** Outdoor walking: mean regressor value in Walking 1 against Walking 2 for each participant; the dashed line is the identity line. Feature content was highly correlated between the two runs (luminance, r = 0.82; sound intensity, r = 0.91; face intensity, r = 0.98).

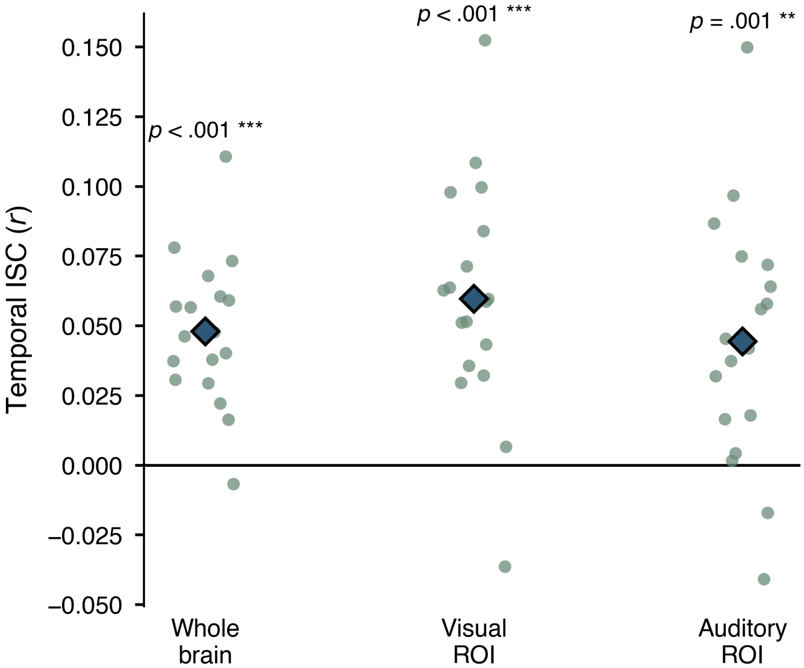


**Supplementary Fig. 3. | Subject-level temporal ISC during movie viewing.** For each participant (n = 18), the HbO time series of each parcel was correlated with the mean time series of the remaining 17 participants (leave-one-out, LOO). Parcel-wise correlations were Fisher z-transformed, averaged within each parcel set and back-transformed to r. Parcel sets were the sensitivity-masked whole brain and the a priori visual (BA17/18/19) and auditory (BA41/42/21/22) ROIs. Each point shows one participant's mean ISC; diamonds show the group mean. Significance was assessed with a two-sided sign-flip permutation test on the Fisher z values (5,000 permutations), and P values were BH-FDR corrected across the three parcel sets. FDR-adjusted P values are shown above each parcel set. Values are reported in Supplementary Table 2.

**Supplementary Table 2 | Temporal inter-subject correlation during movie viewing.** Leave-one-out ISC averaged across participants. p, sign-flip permutation test (5,000 permutations); p (FDR), BH-FDR adjusted across the three parcel sets.

| Parcel set | Mean r | 95% CI | p | p (FDR) |
|---|---|---|---|---|
| Whole brain | 0.048 | [0.036, 0.060] | < 0.001 | < 0.001 |
| Visual ROI | 0.06 | [0.041, 0.078] | < 0.001 | < 0.001 |
| Auditory ROI | 0.044 | [0.025, 0.064] | 0.001 | 0.001 |
